# Assessing the robustness of SNaQ to violations induced by high-level phylogenetic networks

**DOI:** 10.64898/2026.09.01.748505

**Authors:** Joshua Justison, Gustavo A. Ballen, Carlos Acosta-Cortés, Roberto Ceja, Hector Baños, Claudia Solís-Lemus

## Abstract

Phylogenetic networks extend the traditional tree model to capture reticulate evolutionary processes such as gene flow and hybridization. Among available inference tools, SNaQ is a widely used quartet-based method that offers a computationally efficient, statistically grounded approach to network estimation, but is limited to level-1 networks, in which reticulation cycles do not overlap. This assumption is a statistical requirement for identifiability rather than a reflection of biological reality, as many evolutionary scenarios, particularly those involving extensive or closely spaced gene flow, are expected to produce level-2 or higher networks. How SNaQ performs when this assumption is violated remains poorly understood. Here, we systematically evaluate SNaQ’s performance on simulated non-level-1 networks. Because existing network comparison metrics such as hardwired cluster dissimilarity are not true distances beyond level-1, we introduce complementary measures: hybrid cluster compatibility, blob compatibility, and tree-of-blobs comparison, to more directly assess structural recovery. We find that while SNaQ does not recover the exact topology of non-level-1 networks, it reliably infers the circular order of taxa and frequently recovers a tree of blobs compatible with the true network, suggesting the level-1 constraint acts as a form of regularization against overfitting. Recovery of reticulation signal is strongly tied to inheritance proportion and the user-specified maximum number of reticulations, with SNaQ behaving as a conservative estimator that favors strong, well-supported events over finer-scale or overlapping signals. These results clarify the strengths and limits of quartet-based network inference under model misspecification and offer practical guidance for applying SNaQ to complex reticulate histories. Phylogenetic network, gene flow, hybridization, level-1, SNaQ

## 1 Introduction

Reticulate evolutionary processes, such as hybridization, introgression, and lateral gene transfer are increasingly recognized as fundamental to shaping biological diversity across the Tree of Life (Nakhleh, 2011; Ellstrand et al., 1996; Rieseberg et al., 2000; Linder and Rieseberg, 2004; Noor and Feder, 2006; Singh et al., 2022; Lescroart et al., 2023; Pyron et al., 2024; Kong et al., 2025). These complex gene flow processes play vital roles in phenomena ranging from the evolution of virulence in pathogens (Kolb and Brandt, 2020; Arnold et al., 2022) and the adaptation of key traits (Brower, 2013; Lazic et al., 2021), to the broader patterns of species diversification (Meier et al., 2019; Wogan et al., 2023). A comprehensive understanding of evolutionary history, therefore, requires methods that can accurately detect and model these reticulate events.

Although numerous statistical methods have been developed to detect gene flow (Patterson et al., 2012; Pickrell and Pritchard, 2012), these approaches often do not yield an explicit phylogenetic reconstruction of reticulate history. Phylogenetic networks address this limitation by extending the traditional tree framework to incorporate nodes representing reticulation events (Kong et al., 2025; Solís-Lemus, 2026), and have become essential for studying complex species radiations (Edelman et al., 2019; Meier et al., 2019), reconstructing the evolutionary history of domesticated crops (Glémin et al., 2019; Viot and Wendel, 2023; Przelomska et al., 2024), and tracing the evolution of traits (Bastide et al., 2018; Edelman et al., 2019). A range of computational parsimony, distance, likelihood, and Bayesian approaches have been developed for network inference (*e.g.,* Ogilvie et al., 2017; Yan et al., 2021; Solís-Lemus and Ané, 2016; Flouri et al., 2020; Allman et al., 2024b; Holtgrefe et al., 2025b, see also Elworth et al. (2018); Hibbins and Hahn (2022); Kong et al. (2025)), with methods grounded in the Network Multispecies Coalescent (NMSC) (Meng and Kubatko, 2009; Yu et al., 2012) being especially common because they jointly model reticulation (gene flow) and incomplete lineage sorting (ILS) (Maddison, 1997) as sources of gene tree discordance. However, full likelihood and Bayesian NMSC methods are computationally demanding: the combinatorial explosion of possible network topologies as taxa and reticulations increase, together with the analytical complexity of computing gene tree likelihoods conditional on a network, limits their scalability to relatively few taxa and reticulations, restricting their usefulness for many modern genomic studies.

To overcome this scalability limitation, several strategies have been proposed, including distance-based (Willems et al., 2014; Bordewich et al., 2018) and parsimony-based (Kannan and Wheeler, 2012; Van Iersel et al., 2018; Wheeler and Washburn, 2023) methods, as well as divide-and-conquer approaches (Zhu et al., 2019; Kolbow et al., 2025b; Hejase and Liu, 2016). While these accommodate larger datasets, distance- and parsimony-based methods are often statistically inconsistent under biologically realistic scenarios. Distance methods that average evolutionary distances across loci can yield incorrect topologies or unidentifiable parameters (Xu and Ané, 2023), while parsimony-based methods either underestimate reticulations when minimizing their number, or overestimate network complexity when required to display all topologies present in the input gene trees (Mirzaei and Wu, 2015; Bernardini et al., 2024; Zhang et al., 2025), effectively attributing all gene tree discordance to reticulation and failing to account for incomplete lineage sorting or gene tree estimation error. A particularly promising alternative is provided by scalable, model-based summary methods, which achieve computational efficiency while maintaining statistical rigor. These methods typically proceed in two stages: first, individual gene tree histories are inferred from sequence data; second, the distribution of these gene trees is summarized to infer the species network. By decoupling gene tree inference from network inference, summary methods avoid the computationally intensive joint-likelihood calculations while retaining high accuracy. Efficiency is further enhanced by focusing on quartets rather than full gene trees. Specifically, these approaches compute quartet *concordance factors* (CFs), which quantify the proportion of loci supporting each of the three possible unrooted topologies for a given set of four taxa (Solís-Lemus and Ané, 2016; Kong et al., 2024; Allman et al., 2019, 2025a). Because quartets are not independent, a composite likelihood is used to combine quartet likelihoods. The composite likelihood of a candidate network is then evaluated based on the expected CFs under the NMSC model. This framework allows summary methods to scale to genome-wide datasets while providing a statistically principled means of inferring complex reticulate evolutionary histories.

A key limitation of the quartet-based framework is that not all network topologies or parameters are identifiable from CFs alone (Solís-Lemus and Ané, 2016; Baños, 2019; Allman et al., 2024a; Ané et al., 2024; Rhodes et al., 2025). Identifiability is largely restricted to simpler classes of networks, with *level-1* networks being the most thoroughly studied. The level of a network reflects both the number of reticulation events in a given region and the complexity of their arrangement (see Box for a formal definition). Consequently, quartet-based methods are typically designed to explicitly infer networks within the level-1 class. Level-1 networks have long been a popular class for phylogenetic network inference, and substantial progress has been made in their identifiability from sequence data (Gross et al., 2021; Warnow et al., 2025) as well as in their estimation from various substructures, including triplets (Jansson and Sung, 2006), quartets (Keijsper and Pendavingh, 2014; Solís-Lemus and Ané, 2016; Allman et al., 2024a), trinets (Huber et al., 2017), and quarnets (Huber et al., 2018; Holtgrefe et al., 2025b). While some identifiability results exist for a subset of level-2 networks (Englander et al., 2025; Holtgrefe et al., 2025a) or other complex network families (Allman et al., 2025b), no model-based quartet summary methods currently exist to estimate these networks, though a few level-2 constructions have been proposed (Van Iersel et al., 2009). Alternatively, rather than inferring full phylogenetic network, the tree of blobs collapses regions into single nodes to reveal the tree-like structure and is identifiable from quartet concordance factors (Allman et al., 2024b).

Among level-1 network inference methods, SNaQ (Species Networks applying Quartets) (Solís-Lemus and Ané, 2016; Kolbow et al., 2025a) has emerged as a benchmark for scalable, quartet-based approaches. SNaQ infers the level-1 network and associated inheritance probabilities that maximize a composite likelihood function derived from the observed quartet concordance factors. The method employs a heuristic search, using hill-climbing optimization to explore network topology space efficiently. This strategy allows SNaQ to achieve high accuracy while remaining substantially more scalable than full-likelihood approaches, making it well-suited for genome-scale datasets.

#### Box: Definitions

- **Biconnected component (blob):** In graph theory, an undirected graph is considered biconnected if the graph remains connected after the removal of a single vertex. A biconnected component, or blob, is a maximal subgraph where the subgraph remains connected after the removal of a single vertex. Biconnected components with only one node are known as trivial blobs. For the purposes of this study, the blobs we reference are assumed to be non-trivial.
- **level-*k*:** In phylogenetics, a network is considered level-k if any biconnected component on the network has at most k reticulate nodes. Many inference methods are often limited to level-1 networks because these networks were the first shown to be identifiable under the NMSC.
- **Tree of Blobs:** A way of summarizing a phylogenetic network by contracting all the edges within a blob to a single node and suppressing degree-two nodes. The remaining structure is known as the tree of blobs, which represents the treelike portions of the original phylogenetic network. The tree of blobs is identifiable from quartet data under the NMSC and has been used as a part of level-1 network estimation.

However, the level-1 constraint is a statistical requirement for identifiability rather than a reflection of biological reality. Many biologically relevant processes produce networks that are not level-1 (Janssen and Liu, 2021; Justison and Heath, 2023). Level-2 or higher networks are especially common in systems with extensive gene flow or when gene flow is restricted to closely related taxa, as closely spaced reticulation events are more likely to form the same biconnected component. Current level-1 methods, including SNaQ, are incapable of estimating such networks, and it remains unclear how they perform when the underlying network violates the level-1 assumption. Model misspecification can substantially impact phylogenetic inference; for example, assuming tree-like evolution in a reticulate system can produce biased estimates of both branch lengths and topology (Leaché et al., 2014; SolísLemus et al., 2016; Long and Kubatko, 2018; Pang and Zhang, 2022; Dinh and Baños, 2025). This study aims to address this critical gap by systematically evaluating SNaQ under conditions where the true network exceeds level-1 complexity. Our objectives are to characterize potential biases, identify which aspects of the network are reliably estimated under model misspecification, and clarify the limits of quartet-based inference in more complex evolutionary scenarios.

## 2 Methods

### 2.1 Simulating Networks

To systematically evaluate the performance of SNaQ outside of its foundational level-1 assumption, we simulated a diverse set of rooted binary phylogenetic networks using the birth-death-hybridization process implemented in the R package SiPhyNetwork (Justison et al., 2023). We kept the following parameters constant across all simulations: birth rate *λ* = 1.0, death rate *µ* = 0.0, gamma distribution (inheritance proportion) *γ ∼* Beta(10, 10), and the types of hybridization were all equally likely: 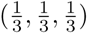 for lineage generative, degenerative, and neutral, respectively. We varied three factors: hybridization rate, number of taxa, and network topology constraint, resulting in 12 distinct simulation settings as shown in Table 1.

**Table 1:**
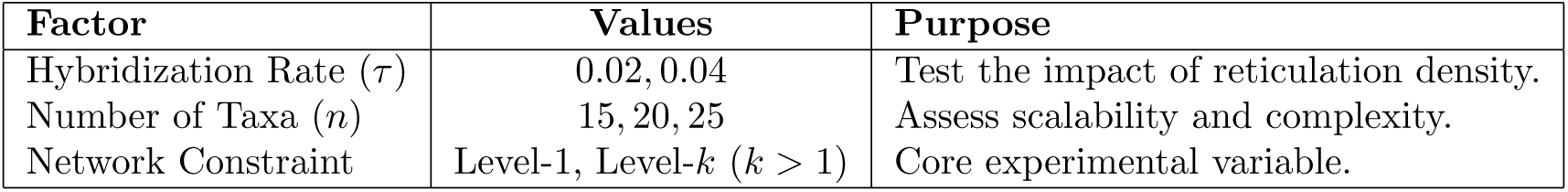
Simulation Settings for Phylogenetic Network Generation.

| Factor | Values | Purpose |
| --- | --- | --- |
| Hybridization Rate ($\tau$) | 0.02, 0.04 | Test the impact of reticulation density. |
| Number of Taxa ($n$) | 15, 20, 25 | Assess scalability and complexity. |
| Network Constraint | Level-1, Level-$k$ ($k > 1$) | Core experimental variable. |

**Table 2:**
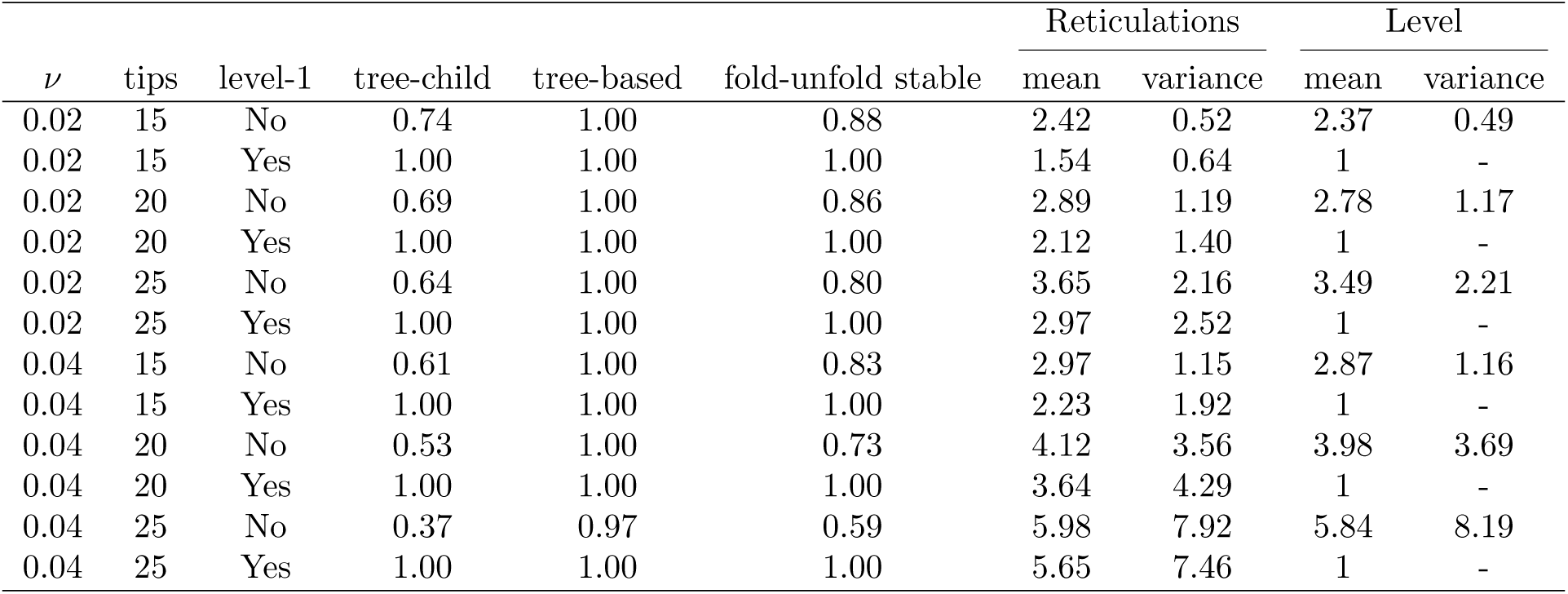
Summary of simulated networks. Values represent the proportion of networks under each simulation condition (rows) with the described properties (columns).

For each of the 12 settings, 150 phylogenetic networks were simulated. To control that unidentifiability is only due to the level of the network, we removed any 2- or 3-cycles (informally, level-1 blobs with 2 and 3 nodes, respectively) from the networks, as these structures are not identifiable in a 4-taxon network from quartet concordance factors (Solís-Lemus and Ané, 2016; Baños, 2019). Additionally, we pruned any edges ancestral to the last stable ancestor (LSA). In a network, the most recent common ancestor of all taxa can be ambiguous, and the LSA is the most recent node that lies on every path from the root to the tips. Edges above the LSA are not identifiable from quartet concordance factors, because they do not carry information about the root (Baños, 2019; Ané et al., 2024).

### 2.2 Simulating Gene Trees

Gene trees were simulated within each phylogenetic network under the network multispecies coalescent (NMSC) model using the Julia package PhyloCoalSimulations (Fogg et al., 2023). To assess the robustness of network inference on sample size, we varied the number of gene trees: 100, 1000, and 10000. For each combination of a single simulated network and a specified number of gene trees, we generated 30 replicate datasets.

The gene trees were summarized into quartet concordance factors (CFs) using the Julia package PhyloNetworks (Solís-Lemus et al., 2017), which measures the proportion of gene trees supporting each possible unrooted quartet topology, serving as the primary input data for SNaQ. Note that because our goal is to assess the identifiability limits of SNaQ, we do not consider gene tree estimation error. Instead, we treat simulated gene trees as known without error, allowing us to isolate network features that remain inestimable even under ideal, error-free conditions.

### 2.3 Phylogenetic Network Inference using SNaQ

We estimated the phylogenetic networks with SNaQ v1.1 (Kolbow et al., 2025a). To establish a high-quality starting point for the network estimation, initial starting trees were estimated with Tree-QMC (Han and Molloy, 2023) under the fast setting. To ensure adequate exploration of network topology space, we performed 45 independent runs for each inference and set the stopping parameter nfail = 600. This setting causes the search to terminate only after 600 consecutive topology proposals fail to improve the log-likelihood. In accordance with common practices for SNaQ inference (Solís-Lemus and Ané, 2016; Tiley et al., 2023), we used an iterative approach when setting the maximum number of reticulations (*h*_max_). The search began by running SNaQ with *h*_max_ = 0, inferring a tree. We then iteratively increased *h*_max_ by 1 up to a predetermined limit, defined as either the number of hybridizations or the number of blobs in the true network, with a maximum *h*_max_ of 5. For *h*_max_ *>* 0, the optimal network estimated at *h*_max_ = *i* was used as the starting point for the subsequent search at *h*_max_ = *i* + 1.

### 2.4 Performance of Network Inference

To evaluate the performance of SNaQ when its level-1 assumption is violated, we used several measures to compare the true network (*N_true_*) and the network estimated by SNaQ (*N_est_*). Given that *N_est_*is constrained to be level-1 while *N_true_* is not, we did not expect topological identity. Instead, our analyses were designed to quantify *how* the estimated network approximates, or fails to approximate, the more complex true history. We assessed performance using metrics for topological distance, accuracy of reticulation inference, and conservation of underlying tree-like and abstract network structures as described below.

#### 2.4.1 Characterizing Simulated Networks

Before analyzing the estimated networks, we first characterized the properties of the true, simulated networks (*N_true_*) for each simulation condition. We computed the distribution of relevant topological features, including the number of reticulations, the number of biconnected components (blobs), the number of nodes in biconnected components, and the network level. We also characterized the parameters governing the coalescent process by summarizing the distribution of internal branch lengths. Finally, to quantify the resulting signal strength, we summarized the difference between observed and expected quartet CFs across all simulated networks, noting the average dominant quartet and the difference between minor as a proxy for the overall level of ILS and reticulation signal.

#### 2.4.2 Concordance Factor Comparison

To assess how well *N_est_* captures the signal in the observed data, we compared three sets of quartet CFs: *CF_obs_*(simulated under *N_true_*, used as input), *CF_true_* (expected under *N_true_*), and *CF_est_* (expected under *N_est_*). For each pair of CF sets, we measured agreement using the average Manhattan distance across all quartets *q ∈ Q*:

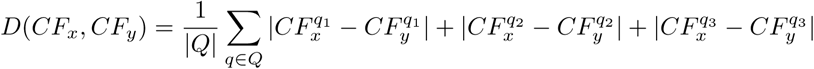

where *CF^qi^* corresponds the proportion of quartets with one of the three unrooted quartet topologies for a quartet *q*. The Manhattan distance corresponds to the total mass differing between two CF distributions. We compute three pairwise distances: *D*(*CF_true_, CF_obs_*) captures the noise introduced by the simulation process (i.e., the discrepancy between perfect and observed CFs under *N_true_*); *D*(*CF_est_, CF_obs_*) measures how well the estimated network fits the data; and *D*(*CF_true_, CF_est_*) measures the residual error between the estimated and true networks, independent of simulation noise.

#### 2.4.3 Quantifying Topological Agreement to Ground Truth

We used two distinct metrics to measure the topological dissimilarity between *N_true_*and *N_est_*. First, we used Hardwired Cluster Dissimilarity (HWCD) which extends the principles of the Robinson-Foulds (RF) distance to phylogenetic networks and is implemented in PhyloNetworks (Solís-Lemus et al., 2017). This metric compares the sets of hardwired clusters (clades) defined by the edges of the two networks. However, it should be noted that this is only a true distance for level-1 networks. Two topologically distinct non-level-1 networks can share the same set of hardwired clusters and thus have a HWCD of 0. The second measure of topological dissimilarity was the Quarnet Consistency Score implemented in SQUIRREL (Holtgrefe et al., 2025b). This score calculates the proportion of the unrooted, four-taxon subnetworks (quarnets) induced by one network that are also induced by the other. This provides a direct measure of how much of the true network’s quartet-level topological information is captured in the estimated network and is known to be a distance metric up to level-2 networks.

#### 2.4.4 Assessing Exact and Partial Recovery of Reticulations

A primary goal was to determine if and how SNaQ captures the true hybridizations present in *N_true_*. First, we assessed the ability of *N_est_* to correctly classify tips as having hybrid ancestry. For every taxon, we recorded its status as either having hybrid ancestry or not in both *N_true_* and *N_est_* and calculated the number of true positives (TP; correctly identified as hybrid), true negatives (TN; correctly identified as non-hybrid), false positives (FP; wrongly identified as hybrid), and false negatives (FN; wrongly identified as non-hybrid), and summarized these values by computing the recall (positive predictive value;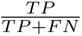) and precision (true positive rate; 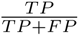).

However, vastly different evolutionary histories may have the same hybrid tips. To better capture whether reticulate events were correctly estimated, we analyzed hybrid cluster compatibility. Each reticulation node in a network defines a hybrid cluster, which is the set of all descendant tips. We compared the set of hybrid clusters from *N_est_* against the set from *N_true_*.

Reticulations between the estimated and true networks were denoted as compatible using two criteria (Figure 2): (i) **exact** when the set of descendant tips in the estimated hybrid cluster was identical to the set in the true hybrid cluster, and (ii) **consistent** when the set of descendant tips in the estimated hybrid cluster was a subset of the tips in the true hybrid cluster. We then call the proportion of all true hybrid clusters that are compatible with a cluster in the estimated network the “hybrid cluster precision”. Similarly, we denote the proportion of all estimated clusters that are compatible with a cluster in the true network as the “hybrid cluster recall”.

**Figure 1:**
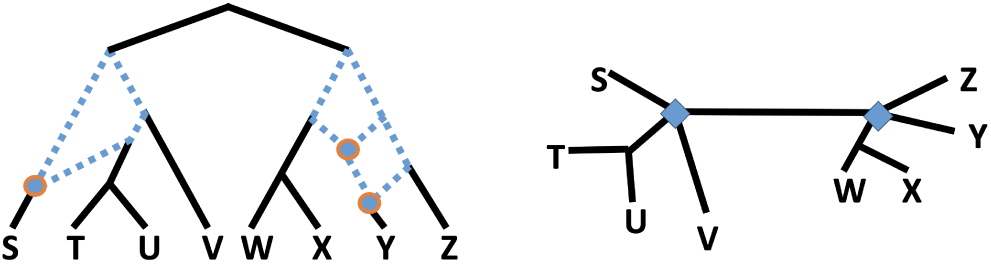
Left: A level-2 network with biconnected components outlined by blue dashed lines. The network is classified as level-2 because the biconnected component on the right contains two hybrid nodes (blue dots with orange outlines). Right: The corresponding unrooted tree of blobs for the network shown on the left. Each biconnected component has been contracted into a single node, represented by blue diamonds. Note that the tree of blobs contains polytomies at each contracted blob.

**Figure 2:**
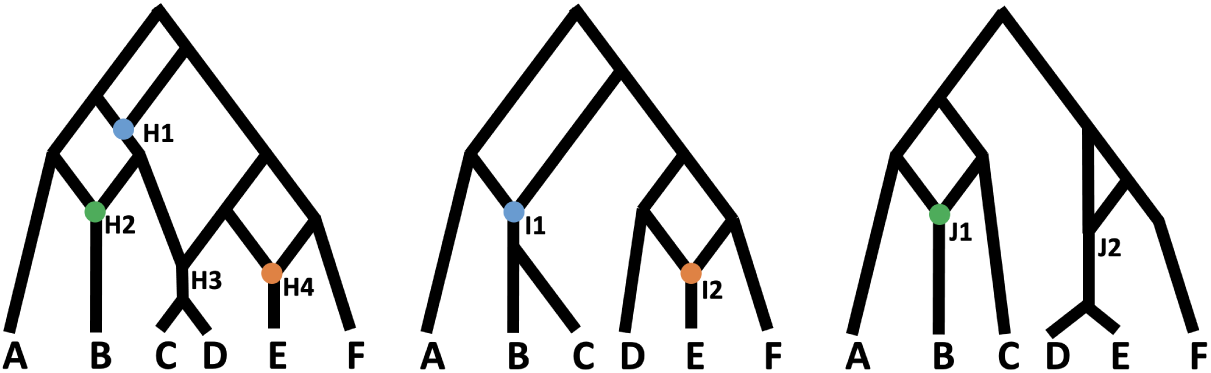
Mapping reticulate nodes between networks: true network (left), estimated network 1 (center), estimated network 2 (right). The reticulate nodes in networks define a set of hybrid descendants. For example, in the true network, hybrid node H1 has *{B, C, D}* as descendants. In estimated network 1, I1 is consistent with H1 because *{B, C}* is a subset of the descendants of H1. Hybrid I2 is consistent with H4 and exact match with H4. In estimated network 2, hybrid J1 is consistent with H1 and H2 while also being an exact match with H2 because the define the same set of descendants (*{B}*). Hybrid J2 has no consistencies or matches.

**Figure 3:**
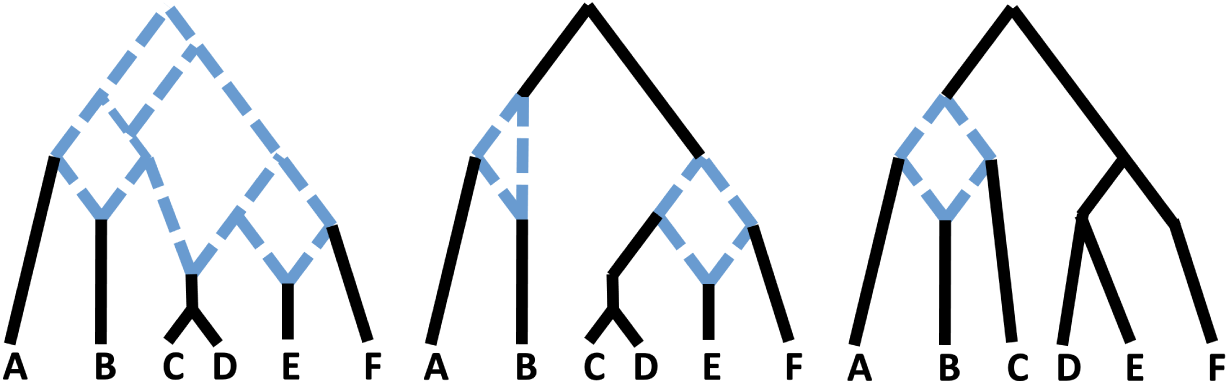
Comparing the blobs in three networks. The true network (left) has one blob that defines the clades *{A}*,*{B}*,*{C, D}*,*{E}*, and *{F }*. The estimated network 1 (center) contains two blobs that are consistent with the true network. In particular, for example, the left blob defines the clades *{A}*, *{B}*, and *{C, D, E, F }*, and every clade in the corresponding blob of the true network is a subset of these clades. Estimated network 2 (right) contains a blob that is inconsistent with the true network. This blob defines the clades *{A}*, *{B}*, *{C}*, and *{D, E, F }*. It is inconsistent because the clade *{C, D}* from the true network is not contained within any of these clades.

We further investigated which properties of a reticulation *h ∈ N_true_* are associated with its recovery in *N_est_*. For each true reticulation (*h ∈ N_true_*), we defined a binary response variable *Y_h_ ∈ {*0, 1*}* based on whether it was found in the estimated network under the exact criterion. We then modeled this response using a set of predictor variables characterizing each reticulation’s properties (each italicized term corresponds to one regressor term): (1) *inheritance proportion* (*γ ∈* [0.5, 1]) associated with the major reticulation edge; (2) *path size*: the number of edges across all paths from the hybrid node to its last stable ancestor; (3) *blob node number* : the number of nodes in the blob containing the hybrid; (4) *blob level* : the number of reticulations in the blob containing the hybrid, and (5) *relative position*: a number of binary variables indicating if the reticulation had *no descendant hybrids in blob* (i.e. an ‘external’ reticulation), *part of another cycle* (i.e., the node lies along one of the paths forming a cycle associated with another reticulation node), or *adjacent to another cycle* (i.e., the paths forming a cycle associated with the reticulate node share one or more edges with the paths that form a cycle associated with another reticulation node, but not the reticulate node itself).

We used two statistical methods in R to identify key characteristics that identify which hybrids were estimated. First, we fit a multivariate logistic regression model on *Y_h_* as the binary response and the predictors listed above using the survival R package (Therneau and Lumley, 2015). to quantify the direction, magnitude, and statistical significance of each property on the probability of detection. Second, we used a decision tree analysis using the caret package (Kuhn et al., 2020) to identify potential non-linear relationships and high-order interactions, and to assess the relative importance of each feature in predicting detection success.

#### 2.4.5 Evaluating Blob-Level Topological Agreement

To evaluate recovery of the tree-like structure of *N_true_* in *N_est_*, we compared their trees of blobs (ToB) (Gusfield et al., 2007), which collapse each blob to a single node and thus capture only the tree-like skeleton of the network. We quantified differences between *ToB*(*N_true_*) and *ToB*(*N_est_*) using two tree distance metrics: the Robinson–Foulds (RF) distance and the Matching Split Information (MSI) distance. While RF measures exact agreement in splits, MSI captures partial agreement between similar splits, making it particularly informative when hybridization events are not recovered exactly, and hybrid clades are split across parental lineages. Because topological differences in ToBs do not necessarily imply incompatible evolutionary histories, we further compared the composition of the blobs themselves. Specifically, a blob in *N_est_* was considered compatible with a blob in *N_true_* if it was either exact (defining the same clades of taxa) or consistent (with each clade in the true blob represented as a subset of a clade in the estimated blob) The proportion of true blobs that were found to be compatible with a blob in the estimated network is denoted as blob precision, while the proportion of estimated blobs that map to a true blob is the blob recall.

#### 2.4.6 Quantifying Topological Agreement on Reduced Networks

We hypothesized that even if *N_est_* is topologically incorrect, it might capture a “reduced” or “canonical” form of *N_true_*. We investigated this by transforming the true and estimated networks into two abstract, stable representations:

### Canonical Networks

Following Pardi and Scornavacca (2015), a canonical network is a unique topological representative of the set of phylogenetic networks that display the same collection of trees. By converting both networks to their canonical forms, we can assess whether they define the same underlying structure while ignoring certain features such as the precise order of reticulation edges within a blob.

### Fold-Unfold Stable Networks

Based on the operations of Huber et al. (2016), the unfold operation *U* (*N* ) converts a network into a multi-labeled tree (MUL-tree) by duplicating the subtrees below reticulation nodes, thereby representing all possible evolutionary paths. The fold operation *F* (*T* ) reverses this process, collapsing isomorphic subtrees to produce a network with a minimal number of reticulations. A network is *fold–unfold stable* if applying these operations in sequence, *F* (*U* (*N* )), returns the original network. By comparing networks through their fold–unfold stable forms, we can determine whether *N_est_* and *N_true_* represent the same set of evolutionary paths, even if their reticulation structures differ.

After transformation, we computed the topological dissimilarity (HWCD) between the corresponding reduced forms. For example, a small dissimilarity between the canonical forms would indicate that *N_est_*captures the core, identifiable structure of *N_true_*, even if the full topology differs. We also explicitly tested if *N_est_* was a subnetwork of *N_true_*, *i.e., N_est_* can be obtained from *N_true_* by removing hybrid edges.

#### 2.4.7 Comparing Displayed Trees

Finally, we compared the sets of bifurcating trees displayed by each network. These trees represent the possible evolutionary histories that a lineage can follow through the network after removing one hybrid edge from each hybridization event. Let *T_true_* be the set of trees displayed by *N_true_* and *T_est_* be the set displayed by *N_est_*.

##### Set Overlap

We first calculate the size of the set intersection (*|T_true_ ∩ T_est_|*) and the set differences (*|T_true_ \ T_est_|* and *|T_est_ \ T_true_|*).

##### Optimal Mapping Distance

Because the set-overlap approach is binary and ignores “near-miss” trees, we also compute an optimal mapping distance. This involves finding a one-to-one mapping between the trees in *T_true_* and *T_est_* that minimizes the total sum of RF distances, providing a more continuous measure of similarity between the two sets of displayed trees.

### 2.5 Assessing the Identification of Circular Order

A blob is said to be *outer labeled planar* (OLP) if it admits a planar embedding in which (1) no edges cross and (2) all taxa lie on the unbounded face. A network is OLP if all of its blobs satisfy these properties. Figure 4 illustrates an OLP network (left), along with two distinct embeddings of a network that is not OLP (center and right). The embedding depicted in the center of such a Figure has edges crossing, while the embedding depicted on the left contains taxa on the bounded face.

**Figure 4:**
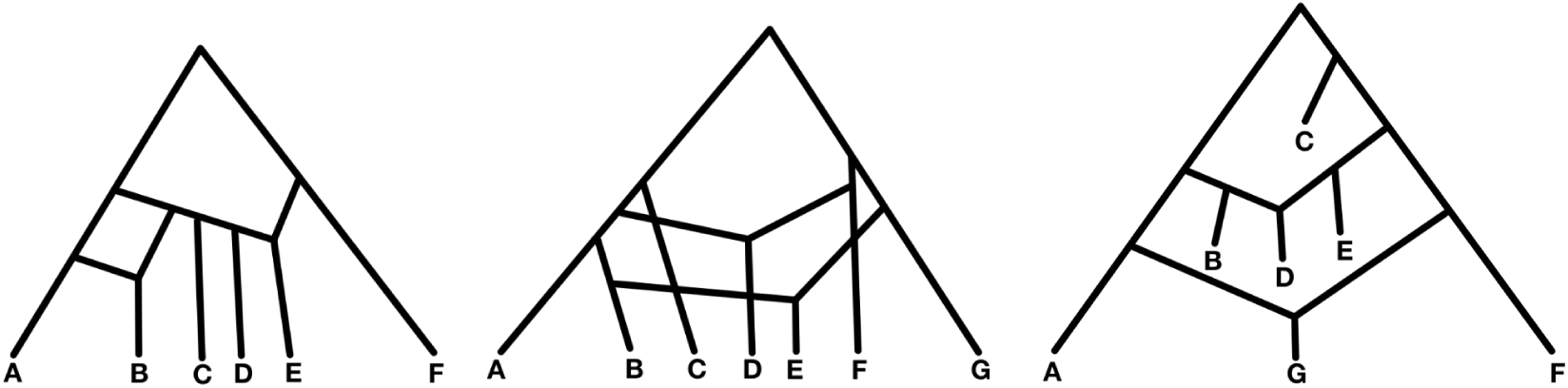
Left: An outer labeled planar (OLP) network. Center: An embedding of a network that is not OLP with edge crossing. Right: An embedding of a network that is not OLP with taxa in a bounded face.

A key feature of OLP networks is that each blob induces a *circular order* of the taxa (up to cyclic permutation), determined by their order of appearance along the unbounded face. For example, the network in Figure 4 (left) induces the circular order (*a, b, c, d, e, f* ). In particular, all level-1 networks are OLP. Moreover, Rhodes et al. (2025) recently showed that, for OLP networks of arbitrary level, this circular order is identifiable from concordance factors. Extending the results in Ceja (2025), we investigated whether SNaQ can recover the circular order under model misspecification. To this end, we simulated data on 187 topologically distinct level-2 rooted networks on six taxa, each containing a single nontrivial 6-blob. We excluded networks with 2-cycles, which, as mentioned before these are not identifiable from concordance factors (Ané et al., 2024), and thus produce data indistinguishable from that of a level-1 network.

For this experiment, for each of the 187 networks, we assigned metric information as follows: (1) all edge lengths were sampled from a gamma distribution with shape and scale equal to 1 (and therefore a mean of 1); and (2) hybridization parameters were sampled from a uniform distribution over the interval (0.2, 0.8). This distribution was chosen to avoid hybridization parameters close to 0 or 1, which would correspond to weak hybridization signals and networks close to level 1. For each network, we simulated gene trees with sample sizes of 200, 500, and 1000.

For each gene tree sample, we ran SNaQ (using default parameters) with *h*_max_ = 1 to estimate a semi-directed level-1 network, *N_est_*. The choice of *h*_max_ was determined to align with the number of blobs in *N_true_*.

We determined the proportion of times that *N_est_* had a circular order consistent with that of *N_true_*. That is, *N_est_* can be embedded in the plane with no edge crossings and all taxa in the unbounded face, with the same circular order as *N_true_*. We also examined the proportion of times that it inferred the correct tree of blobs.

## 3 Results

### 3.1 Characterizing Simulated Networks

We first analyzed the topological properties of the simulated networks (Table 3.1). The simulations yielded a diverse set of network topologies that varied significantly in complexity, driven by the hybridization rate (*ν*) and the number of taxa. For networks that were not conditioned to be level-1, the mode network level was 2 and levels ranged from 2 in the simplest simulation condition (*ν* = 0.02, 15 tips) to 17 in the most complex (*ν* = 0.04, 25 tips); though over 85% of networks had a level less than 6. The variance in both the number of reticulations and network level increased substantially with higher taxon counts and hybridization rates, indicating that the harder simulation conditions produced a varied and complex set of evolutionary histories. When networks were restricted to be level-1, they produced networks with fewer reticulations, indicating a slightly less complex space for network inference aside from the topological restriction of being level-1.

**Table 3:** Logistic regression model summarizing the effects of reticulation properties on hybrid detection success. Hybridization detection is defined as the estimated network having a hybridization that has the exact set of descendants as a hybridization in the true network.

| term | mean estimate | *SE* | *CI<sub>lower</sub>* | *CI<sub>upper</sub>* | *z<sub>score</sub>* | *p<sub>value</sub>* |
| --- | --- | --- | --- | --- | --- | --- |
| Blob Node Number | -0.06 | 0.03 | -0.10 | -0.01 | -2.14 | 3.26e-2 |
| Blob Level | -0.29 | 0.13 | -0.50 | 0.00 | -2.21 | 2.69e-2 |
| Adjacent to another Cycle | -0.39 | 0.14 | -0.68 | -0.14 | -2.74 | 6.14e-3 |
| No Descendant Hybrids in Blob | -0.20 | 0.04 | -0.28 | -0.12 | -4.64 | < 0.0001 |
| Major $\gamma$ | 2.31 | 0.22 | 1.82 | 2.73 | 10.36 | < 0.0001 |
| Other Detected Hybridizations in Blob | 0.15 | 0.01 | 0.14 | 0.16 | 27.32 | < 0.0001 |
| Part of Another Cycle | -1.89 | 0.05 | -1.99 | -1.80 | -35.76 | < 0.0001 |
| Path Size | 0.56 | 0.01 | 0.54 | 0.57 | 67.84 | < 0.0001 |
| $h_{max}$ | 0.28 | 0.00 | 0.27 | 0.29 | 58.23 | < 0.0001 |

Despite their complexity, the vast majority of simulated networks remained tree-based. In fact, only the networks with 25 tips and the highest reticulation rate (*ν* = 0.04) produced non-tree-based networks, suggesting that the simulated networks can be represented as trees with additional hybrid edges (Francis and Steel, 2015). However, other topological properties decreased notably as complexity increased. For example, while 74% of networks were tree-child in the simplest condition (*ν* = 0.02, 15 tips), this proportion dropped to just 37% in the high-complexity scenarios (*ν* = 0.04, 25 tips). In contrast, when restricted to being level-1, by definition networks were always tree-based, tree-child, and fold-unfold stable (Hellmuth et al., 2023). This trend confirms that our dataset captures a difficulty gradient, ranging from simpler networks with certain topological properties to those that are highly reticulated and structurally complex.

The distribution of internal branch lengths of networks was similar across simulation conditions (Figure S1). Across all conditions, the average internal branch length was approximately 0.49. Under the standard multispecies coalescent, for a rooted three-taxon tree this branch length would result in approximately 40% gene tree discordance; this can be seen as a conservative approximation of the amount of discordance we would expect from simulated networks.

### 3.2 Quantifying Topological Agreement to Ground Truth

We measured pairwise distances among *CF_obs_*, *CF_true_*, and *CF_est_* to evaluate how well each network topology fits the simulated data and the true generating process. To quantify topological error introduced by the level-1 constraint, we compared *N_est_* and *N_true_* using two metrics that capture complementary aspects of network structure.

#### 3.2.1 Concordance Factor Comparison

To measure how well the estimated network explains both the underlying evolutionary history and the observed data, we performed a concordance factor comparison.

Initially, we established a baseline for sampling noise by calculating *D*(*CF_true_, CF_obs_*). Firstly, even with a small number of genes (100), the average distance between the expected concordance factors on the true network and the observed concordance factors from simulated data was relatively small ( Fig. 5); for context, if concordance factors were generated at random, the average Manhattan distance would be 0.8. This indicates that even for a modest number of genes, the amount of noise is low and the observed concordance factors provide a reliable approximation of the underlying concordance factors from the true network (though it should be noted that accuracy in CF estimates space does not guarantee topological accuracy, particularly near boundary regions or the center of the simplex where small shifts in CFs can have a disproportionate affect in topological estimation). Around 40% of estimated networks had a smaller distance between the true CFs and expected CFs from the estimated *D*(*CF_est_, CF_obs_*) than the *D*(*CF_true_, CF_obs_*) (Figure 5), indicating that these networks are worse at explaining the underlying expected CFs (from the true network) than would be expected from noise alone. Conversely, when evaluating model fit as the distance between the observed data and the estimated network *D*(*CF_est_, CF_obs_*) the method performed better, with only 35% having a distance worse than the noise in the data *D*(*CF_true_, CF_obs_*) (Figure 6).

**Figure 5:**
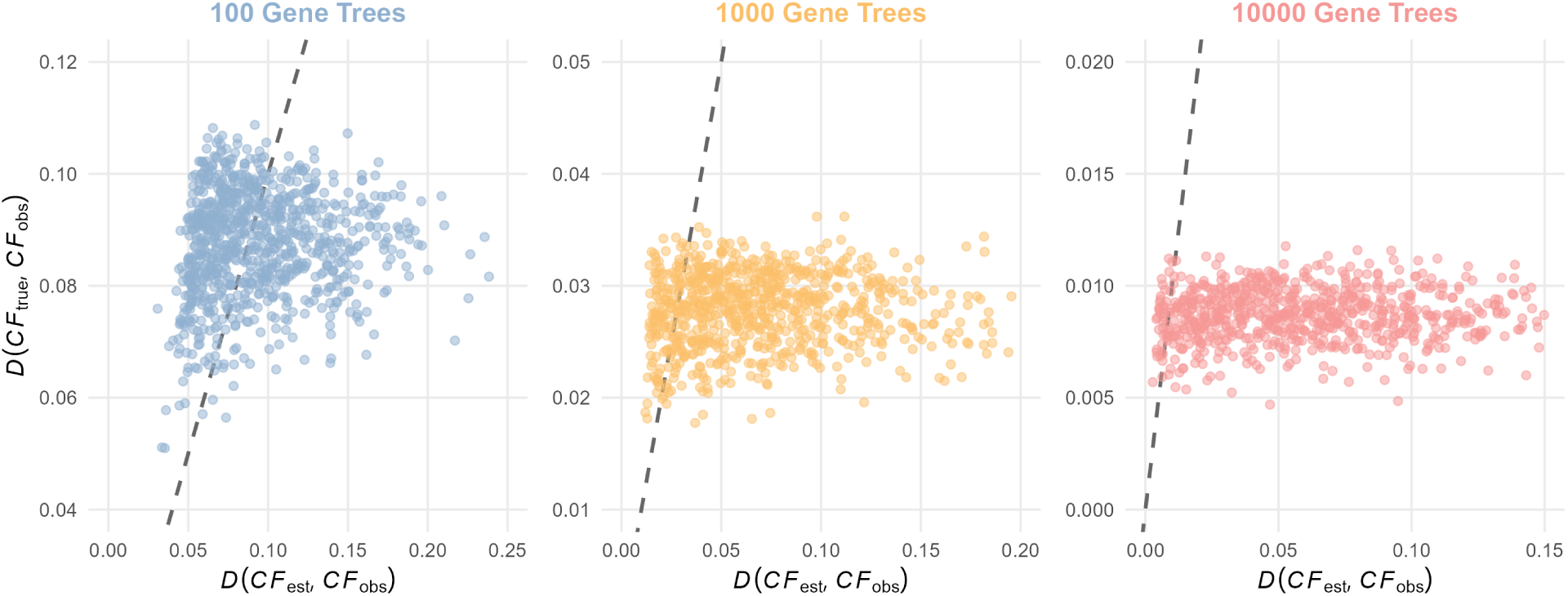
Scatterplot of *D*(*CF_true_, CF_obs_*), the discrepancy between perfect and observed CFs reflecting simulation noise, versus *D*(*CF_est_, CF_true_*), the error in the estimated network relative to the true generating process. Results are aggregated across all tip numbers.

**Figure 6:**
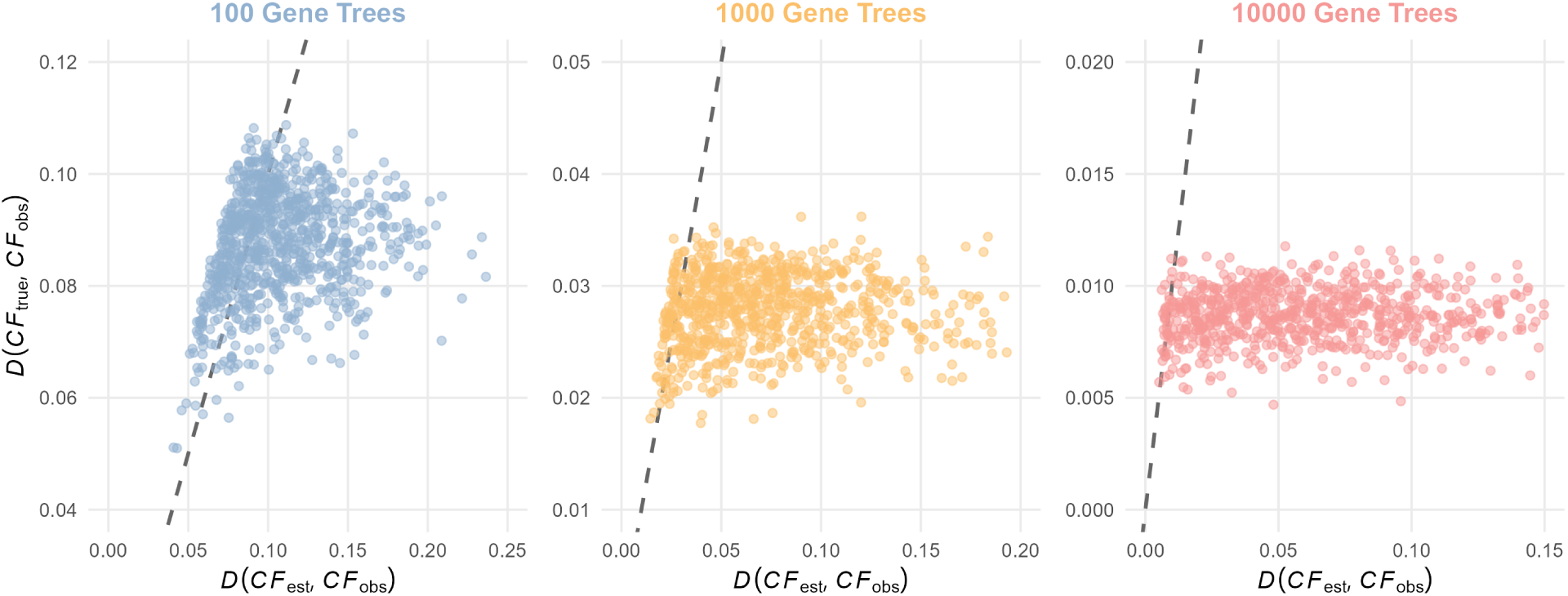
Scatterplot of *D*(*CF_true_, CF_obs_*), the discrepancy between perfect and observed CFs reflecting simulation noise, versus *D*(*CF_est_, CF_obs_*), the discrepancy between the observed CFs and those expected under the estimated network, reflecting how well *N_est_* fits the data. Results are aggregated across all tip numbers.

#### 3.2.2 Hardwired Cluster Dissimilarity

As expected under model misspecification, the hardwired cluster dissimilarity (HWCD), averaged across simulations, increased as the true network level increased (Figure S3), even when *h*_max_ was allowed to grow with the number of reticulations (up to 5; Figure S4). Since SNaQ is restricted to level-1 networks, exact agreement with higher-level true networks is not expected, and HWCD was never zero in those cases. Interestingly, SNaQ rarely recovered a subnetwork of the true network, even when *h*_max_ was set to 1. This could indicate that SNaQ attempts to average or summarize the complex signal reticulate histories as opposed to only returning level-1 subnetworks.

Reducing the true and estimated networks to their canonical or fold-unfold stable forms before computing their HWC dissimilarity led to similar trends as those observed on the full networks (Figure S9). Dissimilarity values for the reduced forms were only modestly lower, reflecting the smaller number of edges (and therefore clusters) after reduction. This similarity in trends indicates that the discrepancies between true and estimated networks persist even after collapsing to these simplified representations, suggesting that the differences are not limited to fine-scale structure but extend to the broader organization captured by the reduced forms.

#### 3.2.3 Quarnet Consistency Score

The quarnet consistency score provides a more rigorous assessment of topological similarity and is a true distance for networks up to level-2. We summarize this score in two ways: (i) the proportion of true quarnets recovered in the estimated network (quarnet precision), which reflects how much of the true signal is captured, and (ii) the proportion of estimated quarnets that are present in the true network (quarnet recall), which reflects how much the estimated network is consistent with the true network.

Quarnet precision remained relatively high across all settings, although the proportion of compatible quarnets decreased as network level increased (Figure 7). Even under model misspecification, SNaQ recovered a substantial fraction of the true signal, capturing approximately 80% of quarnets for level-2 networks, and between 65% to 73% for level-3 networks, depending on the number of gene trees used. This decline with increasing complexity is expected, as higher-level networks contain more quarnets, many of which cannot be represented under the level-1 constraint. Consequently, some true quarnets are necessarily missed, and some inferred quarnets are not present in the true network. Nonetheless, for level-2 networks, quarnet consistency scores indicate that SNaQ recovers the majority of quarnets, demonstrating strong performance even beyond the assumed level-1 class.

**Figure 7:**
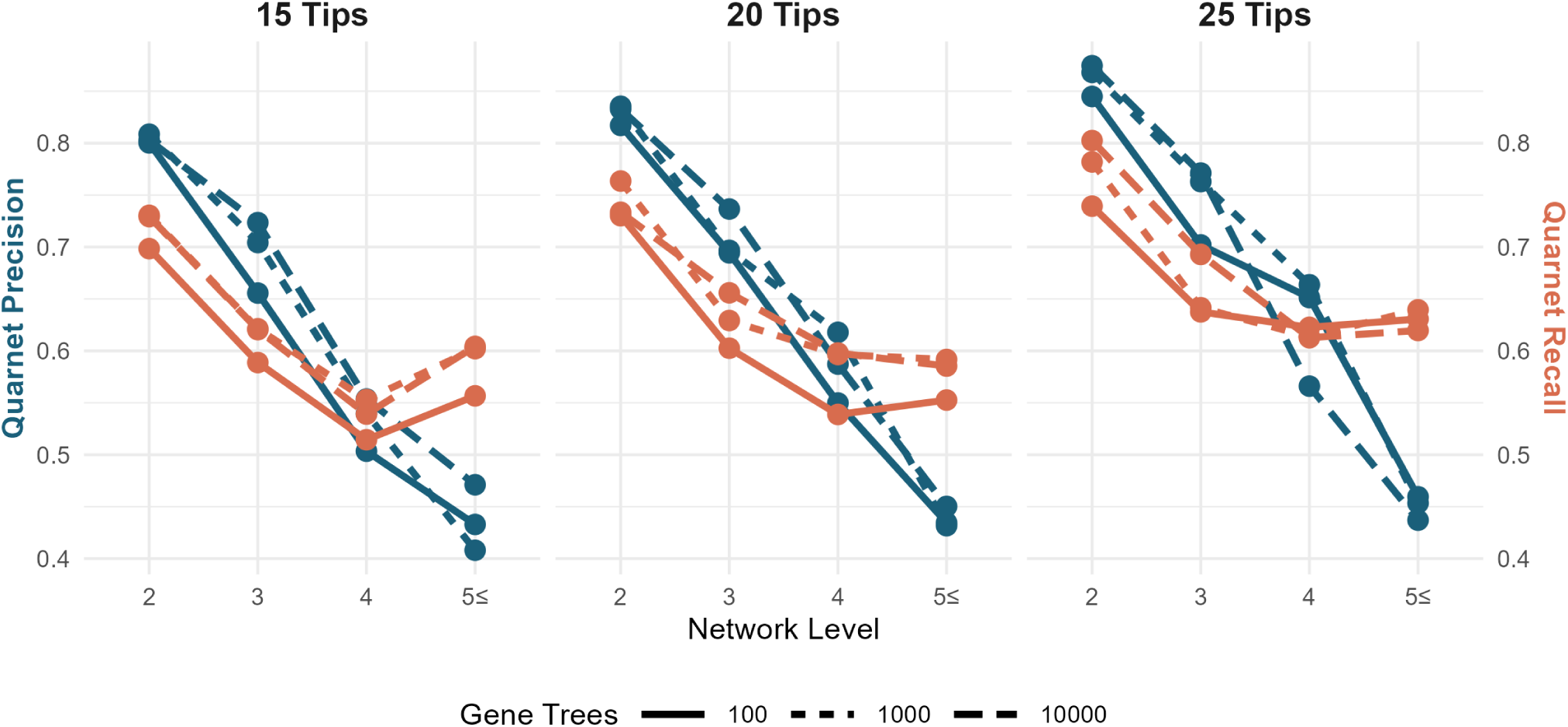
Average quarnet consistency scores when *h*_max_ was set to be the number of blobs in the true network. Each panel aggregates simulations for both values of hybridization rates for 15, 20, and 25 taxa, respectively.

### 3.3 Assessing Exact and Partial Recovery of Reticulations

We evaluated SNaQ’s ability to recover the signal of reticulation at three levels of granularity: the hybrid status of individual tips, the topological placement of hybrid nodes, and the specific properties of reticulations that facilitate their discovery.

#### 3.3.1 Identifying Taxa of Hybrid Origin

At the coarsest level, high precision demonstrated the ability of SNaQ to correctly identify taxa with hybrid ancestry, although low recall indicates that SNaQ is only able to identify a small portion of hybrid individuals on average (Figure 8). Because the level-1 assumption requires each reticulation to involve a distinct set of taxa, allowing larger values of *h*_max_ can force the method to introduce additional, spurious hybridizations. As a result, when we filter results where *h*_max_ matched the number of reticulations in the true network (networks always have a number of reticulations *≥* number of blobs), precision decreased especially for lower-level networks (Figure S5). These results suggest that smaller values of *h*_max_, even when complex hybridization scenarios are expected, may yield more reliable inferences by limiting the inclusion of false hybridization signals.

**Figure 8:**
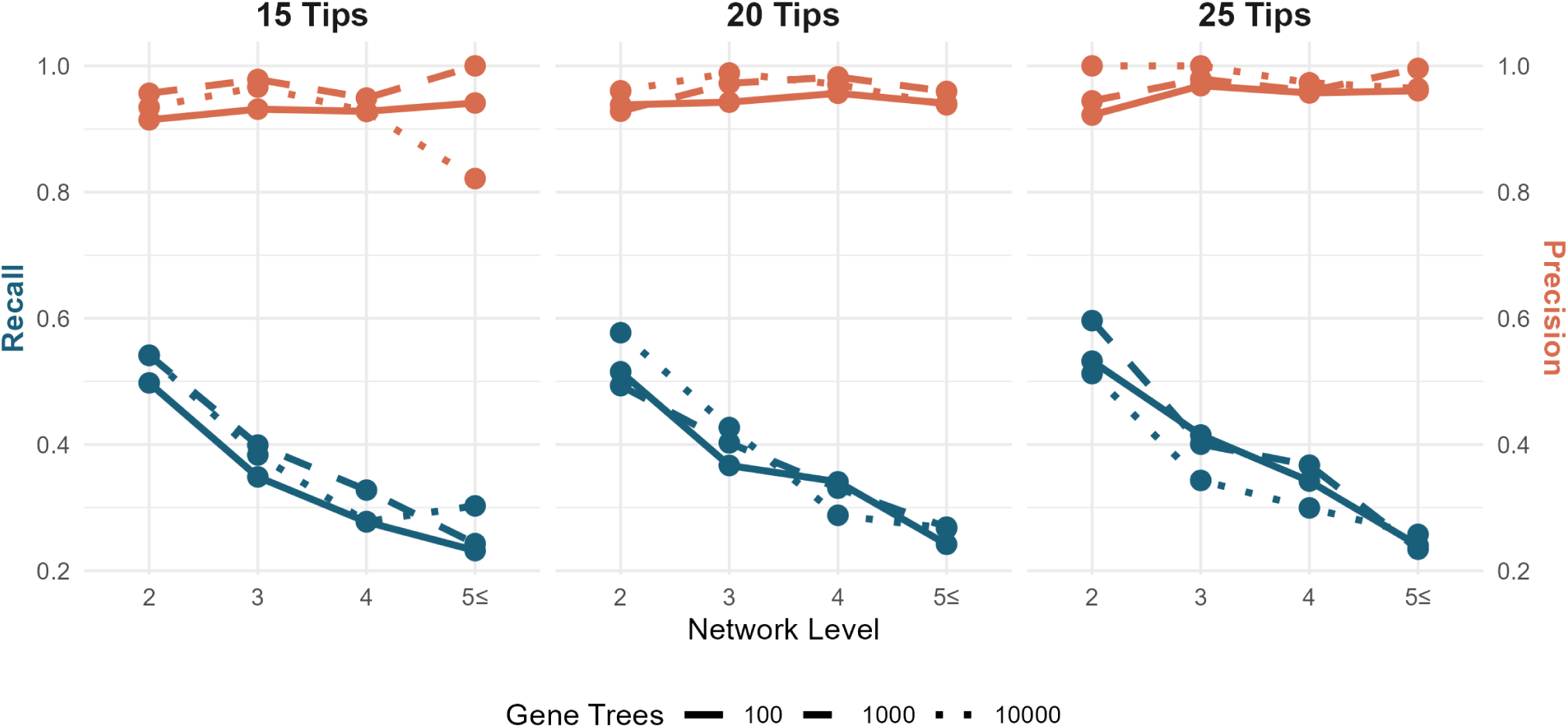
Average recall and precision across replicates for identifying taxa with hybrid descent. The parameter *h*_max_ was set to be the number of blobs in the true network. Each panel summarizes the simulations between the two hybridization rates for 15, 20, and 25 taxa, respectively.

#### 3.3.2 Evaluating Exact and Consistent Hybrid Cluster Compatibility

Hybrid cluster recovery was assessed by comparing the sets of descendant taxa for hybrid nodes. Under the strict exact criterion (requiring exact matching between sets of descendant taxa), hybrid cluster precision was naturally limited by model misspecification, averaging below 20% for level-5 or higher networks (Figure S6). Nonetheless, when using the exact compatibility criterion, hybrid cluster precision remained higher for less complex networks, showing that SNaQ captures many hybrid clades even beyond its level-1 assumptions.

To complement the strict exact criterion, we also evaluated hybrid cluster precision using the consistent criterion, which provides a more generous scope of performance under model misspecification, revealing whether the method captures any true signal and whether its inferred hybrids are generally reliable. In particular, we focus on two metrics that consider subsets of hybrid clades. First, we measured the proportion of true hybrid clade subsets recovered in the estimated networks, which reflects how often SNaQ captures part of a true hybrid clade even if the full set of descendants cannot be perfectly grouped (Figure 9 left). For level-2 networks, approximately 50% of true hybrid clade subsets were recovered in the estimated networks, showing that even partial clades are frequently identified. Second, we measured the hybrid cluster recall, indicating how often inferred hybrids correspond to true reticulations. Across all network levels, including level-5 or higher, over 80% of inferred hybrid clade subsets corresponded to true hybrid clades (Figure 9 right), indicating that when SNaQ predicts a hybrid, it is usually meaningful. Together, these results highlight that SNaQ captures much of the correct phylogenetic signal in terms of identifying hybrid clades. It may underestimate full clade membership due to level-1 constraints, but its inferred hybridizations generally reflect true signal of gene flow.

**Figure 9:**
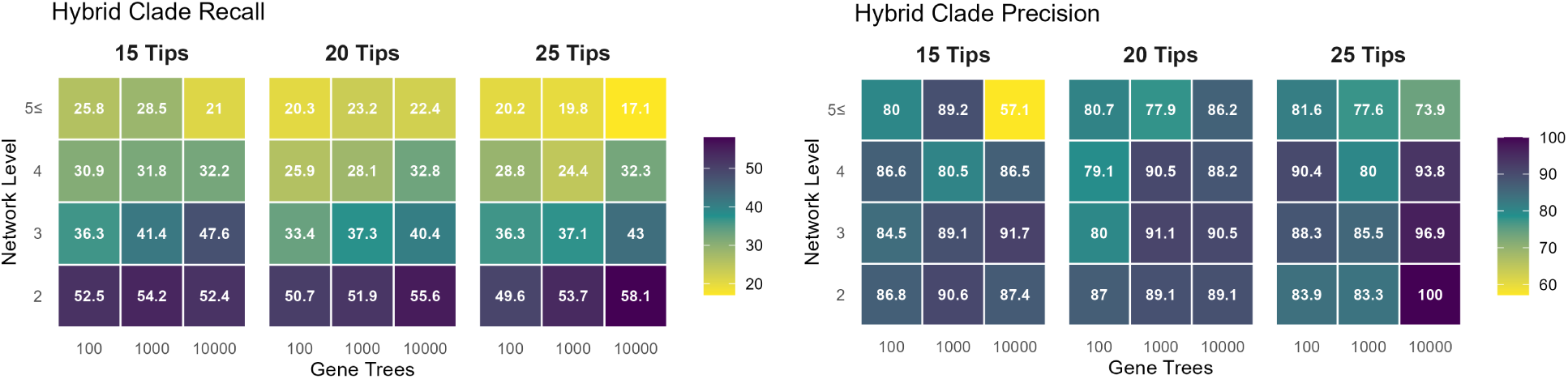
Left: Percentage of true hybrid clades that are consistent in the estimated network (recall). A hybrid clade is considered consistent if the estimated network contains a hybrid node with a subset of the true clade’s descendants, regardless of whether the exact topological relationships are preserved. Right: Percentage of estimated hybrid clades that broadly correspond to a true hybrid clade (precision).

#### 3.3.3 Identifying Properties of Recovered Reticulations

To understand the specific properties and types of reticulation events that SNaQ can recover, we analyzed the characteristics of true reticulations. Logistic regression revealed that detecting other hybridizations in the same blob as a hybridization severely limits recovery, whereas the signal for hybridization itself (represented by *γ*) has a strong positive effect on the recovery of the event (Table 3). These results indicate that for complex series of reticulations where many events are contained in the same blob, only a select few can be detected, likely those with a strong reticulation signal (*γ* values near 0.5). Additionally, general blob complexity measures such as total blob size and the number of reticulations within the blob affected hybridization detection. Interestingly, other reticulation-specific features, including path size, whether the hybridization was external, part of another hybrid cycle, or had adjacent cycles sharing edges, had a comparatively weaker effect on detection (Table 3 and Table 4). The choice of the maximum number of reticulations allowed (*h_max_*) also played a significant role. Higher *h_max_* values naturally increased the probability of recovering any given true reticulation; however, this increased sensitivity comes at the potential cost of topological accuracy elsewhere in the network. In conclusion, with the notable exception of the inheritance proportion (*γ*), the broader context of the hybridization in relation to others within its blob appears to matter more for successful detection than the specific topological qualities of the reticulation itself.

**Table 4:** Feature importance ranking from random forest classification evaluating predictors of hybridization detection success. Importance values represent the mean relative variable importance across permutations, normalized relative to the top predictor (100%), alongside standard deviations (*SD_importance_*) and non-parametric bootstrap 95% confidence intervals (*CI_lower_* and *CI_upper_*).

| term | Mean Importance | *SD<sub>importance</sub>* | *CI<sub>lower</sub>* | *CI<sub>upper</sub>* |
| --- | --- | --- | --- | --- |
| Other Detected Hybridizations in Blob | 100.00 | 0.00 | 100.00 | 100.00 |
| Blob Size | 64.68 | 2.01 | 61.15 | 68.58 |
| Path Size | 58.89 | 4.43 | 53.05 | 69.72 |
| Blob Level | 56.82 | 2.82 | 52.78 | 62.62 |
| $h_{max}$ | 34.65 | 2.20 | 30.68 | 38.39 |
| Major $\gamma$ | 34.07 | 1.35 | 31.73 | 36.76 |
| Part of Another Cycle | 21.36 | 1.59 | 18.17 | 24.15 |
| No descendant Hybrids in Blob | 3.12 | 1.21 | 1.10 | 5.64 |
| Adjacent to another Cycle | 0.00 | 0.00 | 0.00 | 0.00 |

### 3.4 Evaluating Tree of Blobs Topological Agreement

To assess SNaQ’s ability to estimate the treelike segments of an evolutionary history, we assessed the recovery of the Tree of Blobs (ToB) using both strict distance metrics and compatibility measures.

#### 3.4.1 Assessing Topological Error in Trees of Blobs

Robinson-Foulds (RF) distances between the true and estimated ToBs indicated relatively low discordance, reflecting that the overall treelike portion of the network topology was often well recovered (Figure 10). Matching Split Information (MSI) distances were consistently lower than RF distances, possibly suggesting that many of the topological differences were local or due to subtle rearrangements around underestimated reticulations (see Discussion). For example, when SNaQ failed to infer a reticulation, the associated hybrid clade was sometimes split between its two parental lineages in the estimated ToB. While this generates a large penalty under the RF metric, the lower MSI scores indicate that the major phylogenetic splits were preserved. These results show that, even when some reticulations are missed, SNaQ accurately recovers the backbone tree structure and the dominant evolutionary relationships among taxa. Additionally, MSI distances actually decreased as network complexity (level) increased; possibly resulting from the tree of blobs (ToB) becoming increasingly star-like as a larger proportion of the network collapses into a single giant blob. Star-like ToBs inherently would have fewer splits to match between ToBs, indicating less information as a whole.

**Figure 10:**
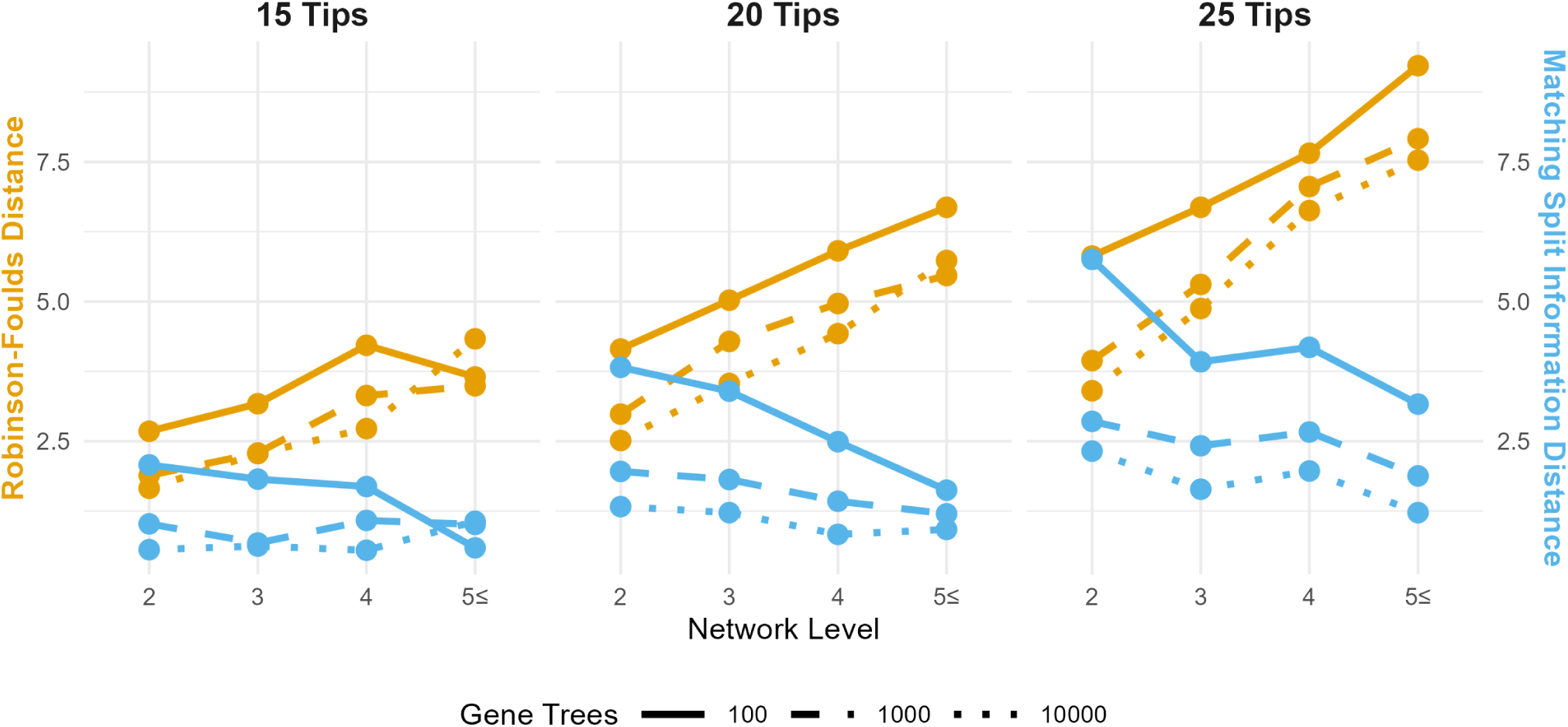
Average distances between the ToB of the true network and the ToB of the estimated network. Orange denotes the Robinson-Foulds distance, while blue denotes the split information distance.

#### 3.4.2 Assessing Exact and Compatible Blob Recovery

In line with the low distances observed between trees of blobs, SNaQ had a high blob precision (the proportion of true blobs that were consistent with a blob on the estimated network). Across all network levels, over 70% of true blobs were recovered in the estimated networks (Figure 11), and most inferred blobs were consistent with those in the true network (Figure S8), leading to high levels of blob recall. These results indicate that, even when complex reticulation patterns are simplified under the level-1 constraint, the inferred blobs largely reflect the true evolutionary relationships. In many cases, this simplification occurs through the decomposition of larger, more complex blobs into smaller structures that remain consistent with the true network. When *h*_max_ was set to match the number of reticulations (rather than the number of blobs), the proportion of compatible blobs decreased (Figure S7), suggesting that additional inferred reticulations can fragment true blobs into smaller, less consistent units.

**Figure 11:**
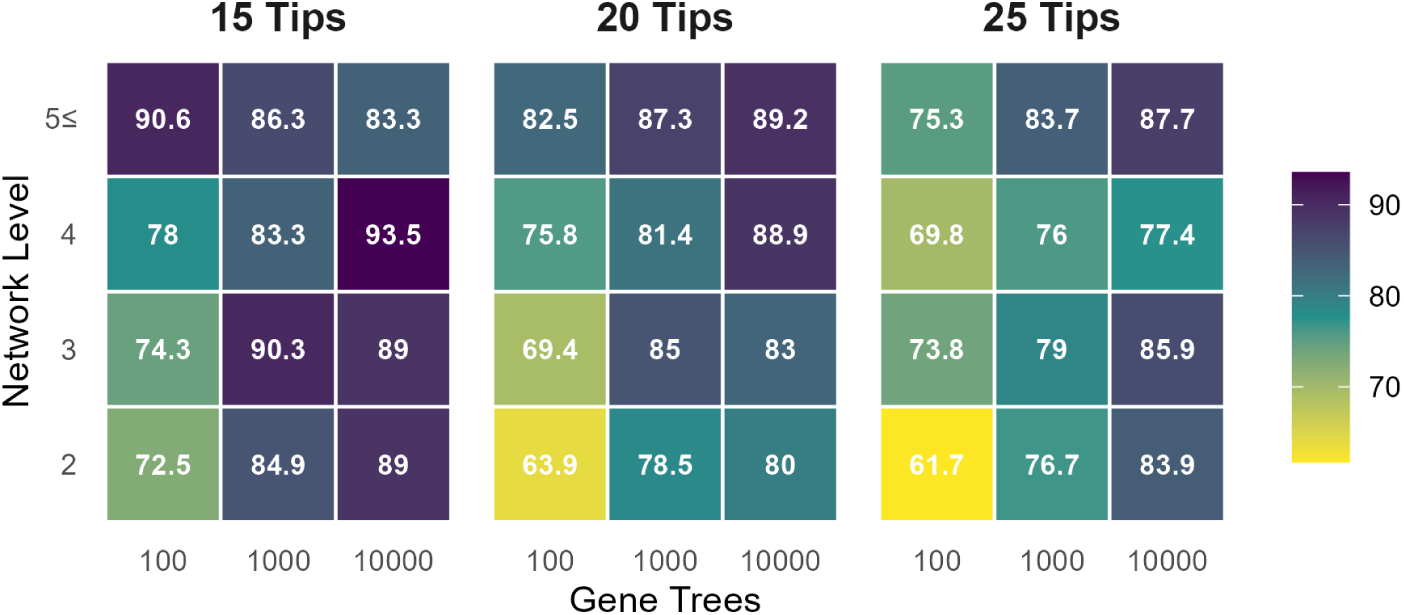
The average proportion of blobs on the true network that were found on estimated networks when the number of hybridizations is set as the number of true blobs.

### 3.5 Comparing Displayed Trees

Increasing network complexity (higher level and reticulation number) led to greater topological discordance between true and estimated displayed trees A network with *h* reticulations has at most 2*^h^* displayed trees, leading to a combinatorial explosion of total paired RF distances across all displayed trees (Figure S10). However, even when controlling for the number of displayed trees by computing average paired RF distance, topological error rose steadily with network level across all dataset sizes (Figure S11). For example, in networks with 15 taxa, the average paired RF distance increased from approximately 4.5 at Level 2 to over 9.0 at levels greater than 5, this represents that nearly 40% of internal splits are not shared between paired trees. Increasing the number of sampled gene trees consistently reduced estimation error.

### 3.6 Assessing the Identification of Circular Order

Table 5 shows the proportion of times SNaQ successfully recovered the circular order and the tree of blobs for all 186 level-2 OLP networks on 6 taxa. While SNaQ did not always recover the correct tree of blobs, it consistently recovered a network with a circular order that was consistent with that of the network used to generate the data. Therefore, SNaQ seems to be very robust in the inference of the circular order, for higher-level networks.

**Table 5:** Proportion of times SNaQ successfully recovered the circular order and tree of blobs for all 187 rooted OLP bloblet networks on 6 taxa with no 2-cycles, across gene tree sample sizes *g* = 200, 500, 1000.

| Feature | $g = 200$ | $g = 500$ | $g = 1000$ |
| --- | --- | --- | --- |
| Circular order | 100% | 96% | 96% |
| Tree of blobs | 66% | 66% | 70% |

## 4 Discussion

Our goal was to assess the performance of SNaQ when its key assumption of a level-1 underlying network is violated. Assessing performance, however, requires comparing inferred and true networks, a task that remains challenging due to the lack of standardized and widely accepted metrics for network comparison. While the Hardwired Cluster Dissimilarity (HWCD) is widely used, its use is limited by the fact that it is only a true distance metric for level-1 networks, as distinct higher-level networks may share identical hardwired clusters and thus yield a score of zero. To address this limitation, we developed several complementary measures: hybrid cluster compatibility, blob compatibility, and network reduction comparisons, to more directly assess structural similarity and quantify SNaQ’s ability to recover reticulate signal. While these measures provide useful insights, their application highlights the need for more robust and principled tools for network comparison.

Hybrid cluster compatibility was designed to identify estimated clusters that capture the signal of true hybridization, but in practice it can be both too lenient and too strict. It is lenient because it considers only the descendants of a reticulation while ignoring the parental lineages. In many applications, the donor lineages contributing to gene flow are of primary interest, yet distinct reticulate histories can produce hybrid nodes with identical descendant sets. This issue is exacerbated in cases of stacked hybridization, where ignoring donor lineages obscures which specific signal has been recovered. Additionally, the measure does not account for relationships within the cluster. Conversely, as a binary measure, it can be overly strict: any inclusion of incorrect taxa results in a score of zero, regardless of the degree of overlap. A more graded, distance-based formulation (e.g., based on symmetric differences of taxon sets) could provide a more informative assessment. Notably, however, our results on the tree of blobs suggest that SNaQ more reliably recovers tree-like relationships.

Blob compatibility exhibits similar limitations. In particular, it strongly penalizes incorrect signal and may therefore underestimate the recovery of true reticulate events. This is especially relevant when multiple hybridization events occur within a single true blob: if the estimated network resolves these into separate blobs, the metric may classify the result as incompatible. Such patterns may arise under the NMSC when lineages descending from a reticulation fail to coalesce and instead traverse different parental paths (Zhu et al., 2016), giving rise to “weakly displayed” subnetworks. Current metrics lack a theoretical framework to account for these fragmented yet partially correct reconstructions. As with cluster compatibility, blob compatibility also ignores internal relationships within clades, focusing solely on membership. Overall, while substantial work remains in developing principled metrics for comparing phylogenetic networks, the measures introduced here provide initial steps toward more fine-grained and interpretable comparisons of network structure.

Our systematic evaluation of SNaQ highlights both the complexity of phylogenetic networks and the impact of violating the level-1 assumption on inference. As expected, the method does not recover the exact topology of non-level-1 networks, resulting in substantial topological discrepancies. However, SNaQ can still provide useful information by recovering a compatible tree of blobs (ToB) structure and (to a lesser extent) the displayed trees, even when the full network contains features whose identifiability is not well understood. By constraining inference to level-1 networks, SNaQ effectively simplifies complex, overlapping reticulation cycles that it cannot represent, collapsing them into simpler structures or treating them as discordance.

For practitioners, this suggests that the tree-like components of the signal inferred by SNaQ likely remain informative, even when the inferred reticulation edges and within-blob structure differ from the true evolutionary history. In this sense, the level-1 constraint, while restrictive, may act as a form of regularization that prevents overfitting and preserves dominant phylogenetic signal. Consistent with this interpretation, we find that certain topological features are reliably recovered despite overall model misspecification: in particular, the circular order is almost always inferred correctly, and the estimated tree of blobs is frequently compatible with the true network. This observation aligns with prior work showing that methods targeting the ToB can yield results consistent with SNaQ (Allman et al., 2025a).

Because SNaQ is restricted to the level-1 model, it cannot fully reconstruct evolutionary histories in which biconnected components (blobs) contain multiple, overlapping reticulations. When reticulation events are well separated by identifiable tree-like edges, SNaQ can often recover a subset of these events within a blob. In contrast, in highly reticulate clades—particularly those with stacked or non-galled reticulations—it struggles to recover the true evolutionary signal. Such configurations produce overlapping clusters of descendant taxa, which are difficult to represent within a level-1 framework where reticulation cycles must be disjoint.

As a consequence, in attempting to fit the data under this constraint, SNaQ may shift parental lineage assignments or exclude subsets of descendant taxa from inferred hybrid nodes. This behavior explains why the method often recovers consistent clusters that reflect subsets of the true signal, yet fails to reconstruct the exact hybrid clades. More broadly, it suggests that inferred reticulation edges in SNaQ should not always be interpreted as discrete biological events. Instead, they may represent aggregated or simplified signals of gene flow. This interpretation is supported by the observed correlation between inheritance proportion (*γ*) and detectability: SNaQ reliably identifies strong gene flow signals, while weaker or overlapping signals are likely merged into dominant events or absorbed as incomplete lineage sorting. In complex scenarios, an inferred level-1 reticulation may therefore be best understood as a proxy for a broader “zone” of reticulation rather than a single event.

The accuracy of inference was strongly influenced by the maximum number of reticulations (*h*_max_) permitted during the search. In general, constraining the model to fewer reticulations produced a less erroneous signal, albeit at the cost of recovering a smaller fraction of the total reticulate history. This pattern suggests that SNaQ can act as a conservative estimator, preferentially identifying the strongest and most well-supported reticulation events while overlooking finer-scale or overlapping signals. Given the risk of overfitting, selecting an appropriate upper bound on the number of reticulations is therefore critical. Complementary analyses aimed at detecting gene flow can help guide this choice. In particular, summary statistic approaches such as the *D*-statistic (ABBA–BABA test) (Green et al., 2010), QuIBL (Edelman et al., 2019), F3 (Reich et al., 2009), MSCquartets (Rhodes et al., 2020), HyDe (Blischak et al., 2018), and likelihood-based tests provide a robust framework for identifying signals of introgression. These methods can help establish an upper bound on the number of reticulations present, although, depending on the structure of gene flow, SNaQ may only reliably recover a subset of these events. Another reasonable way to estimate the number of reticulations is by inferring first the tree of blobs of a network. The tree of blobs is known to be identifiable from quartet concordance factors (Allman et al., 2024b; Rhodes et al., 2025), and recently several methods for its inference have been proposed, including TINNiK (Allman et al., 2024b), TOB-QMC (Dai et al., 2026), and ECToBlob (Rhodes et al., 2026). Such methods can provide an estimate by using the number of blobs as a proxy, where blobs correspond to multifurcations in the inferred tree.

It is important to note that any quartet-based network inference method is susceptible to anomalous networks in which the most frequent gene tree is not present in the true network (Ané et al., 2024). Inferring a network from data generated by an anomalous network is therefore highly likely to produce erroneous taxon relationships. As explored in Ané et al. (2024), although anomalous networks are rare, they do exist, and when finite-sample error is taken into account, even networks that are not anomalous in theory may behave like such in empirical data.

## 5 Conclusions

Given these limitations, we advise caution when interpreting the fine-scale details of networks estimated by level-1 methods. We propose three key guidelines for interpretation:

1. **Underlying treelike structure:** The relationships defined by the underlying treelike structures of the displayed trees and ToB appear robust to model violations, even when reticulation estimates are uncertain.
2. **Reticulation as a flag:** Inferred reticulation edges should be treated as hypothesis generators rather than definitive histories. A detected edge indicates strong evidence for gene flow involving that specific clade, but the precise donor-recipient relationship may be more complex (e.g., involving ancestral relatives or multiple overlapping events) than the level-1 graph depicts.
3. **Indicators of complexity:** Networks may be well estimated by SNaQ if they are indeed level-1; however, it is rarely known *a priori* if the underlying phylogeny satisfies this constraint. As such, any network estimated with SNaQ should be paired with a goodness-of-fit test to assess whether the observed concordance factors can reasonably be explained by the estimated topology. If the concordance factors are poorly explained, this may indicate a more complex reticulate history (or another model violation). Additionally, if the diversification dynamics of the system are known, the results could be paired with a simulation study to assess the probability of generating level-1 networks under the assumed evolutionary model (Justison and Heath, 2023).

The true evolutionary history of many groups is undoubtedly more complex than a level-1 network. While SNaQ and other level-1 approaches will incorrectly estimate the topology in these cases, they have utility in recovering major signals in the reticulate history and remain some of the most scalable approaches for phylogenetic networks when used with caution.

Current efforts towards highly scalable, model-based network estimation are promising (Zhu et al., 2019; Kolbow et al., 2025b). These divide-and-conquer approaches focus on first estimating networks on smaller subsets of taxa and then amalgamating the results into a single, larger network. Notably, InPhyNet (Kolbow et al., 2025b) is able to quickly estimate networks with thousands of taxa, while PhyloNet (Zhu et al., 2019) applies no level-1 constraints. Interestingly, SQUIRREL (Holtgrefe et al., 2025b) also first infers quarnets (four-taxon networks) and uses these to construct a parsimonious network, reinforcing the utility of quartet-based modularity. Similarly, NANUQ+ (Allman et al., 2025a) and NetCS (Dai and Molloy, 2026) employ a divide-and-conquer strategy by using the tree of blobs and resolving each multifunction into a level-1 cycle using quartet data.

Alternatively, recent identifiability results may enable the estimation of networks beyond level-1. Notably, level-2 networks have been shown to be encoded by their quarnets and identifiable from sequence data (Englander et al., 2025). Additionally, it has been shown that tree-child galled networks are identifiable from quartet concordance factors (Allman et al., 2025b), allowing method developers to leverage existing methodology on quartet concordance by extending the search space to include these more complex classes.

Phylogenetic network space is complex, and there are many different classes of networks that denote specific topological properties (see Kong et al., 2022). Many of these classes have ideal theoretical properties and computationally efficient algorithms that may aid in network construction as researchers continue to develop more scalable and accurate inference methods. As we move toward methods capable of handling more general networks with many taxa, these level-1 approximations remain valuable summaries of complex biological reality.

## Code Availability

All simulation and analysis code used in this study is publicly available at https://github.com/solislemuslab/network-simulation-study under the MIT license.

Additionally simulated data and scripts are avaliable at: http://datadryad.org/share/LINK_NOT_FOR_PUBLICATION/xG3VHus7aNFgm64Ag2IRCD2SB1KhwwWzkcud6JzW3Mo

## Acknowledgements

This work was partially supported by the National Science Foundation (DEB-2144367 to C.S.L.; DMS-2331660 to H.B.). G.A.B. was supported through a postdoctoral fellowship by FAPESP (#2023/07838-1).

---

**Figure S1:**
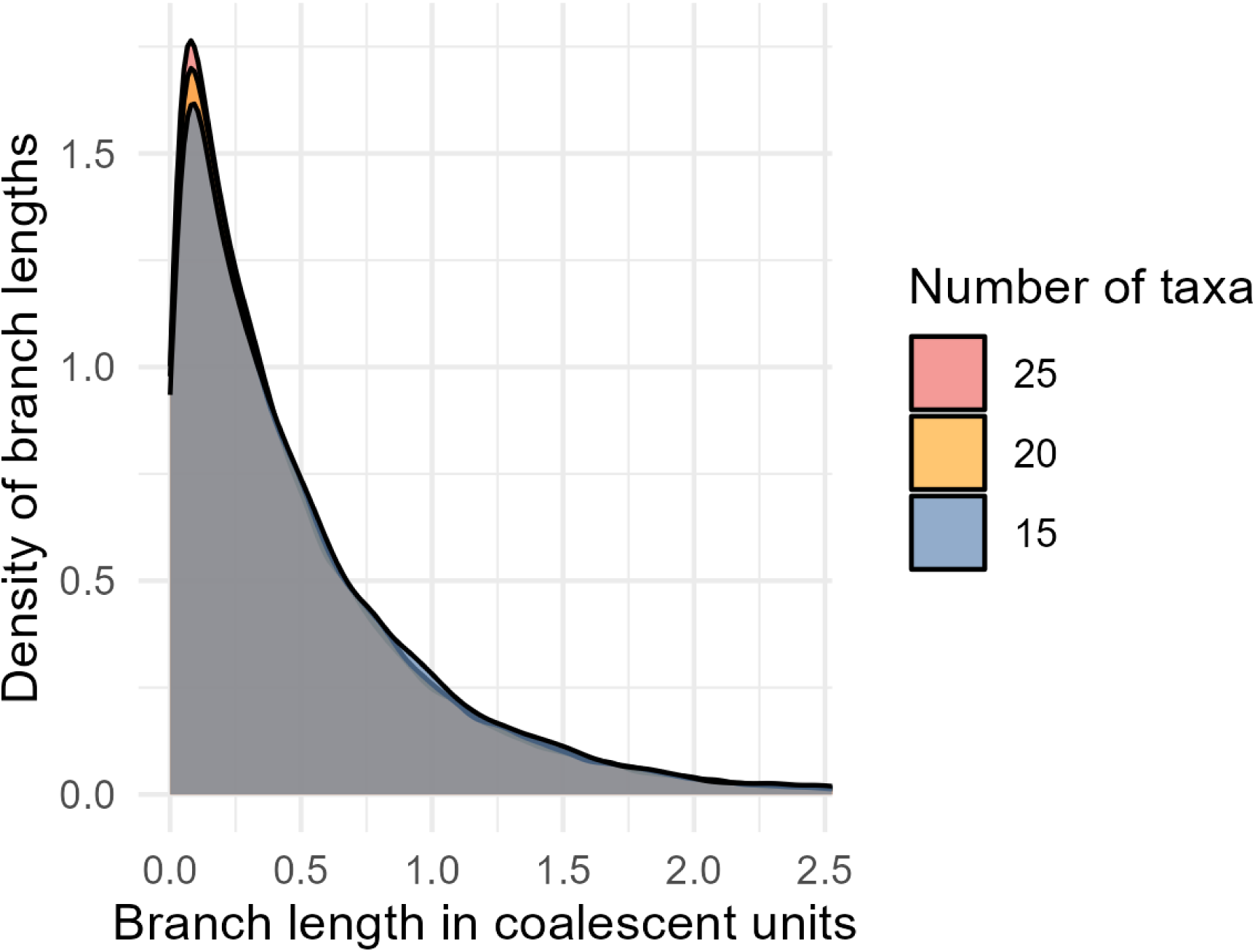
Distribution of internal branch lengths across simulations conditioned to be greater than level-1

## A Supplementary Material

**Figure S2:**
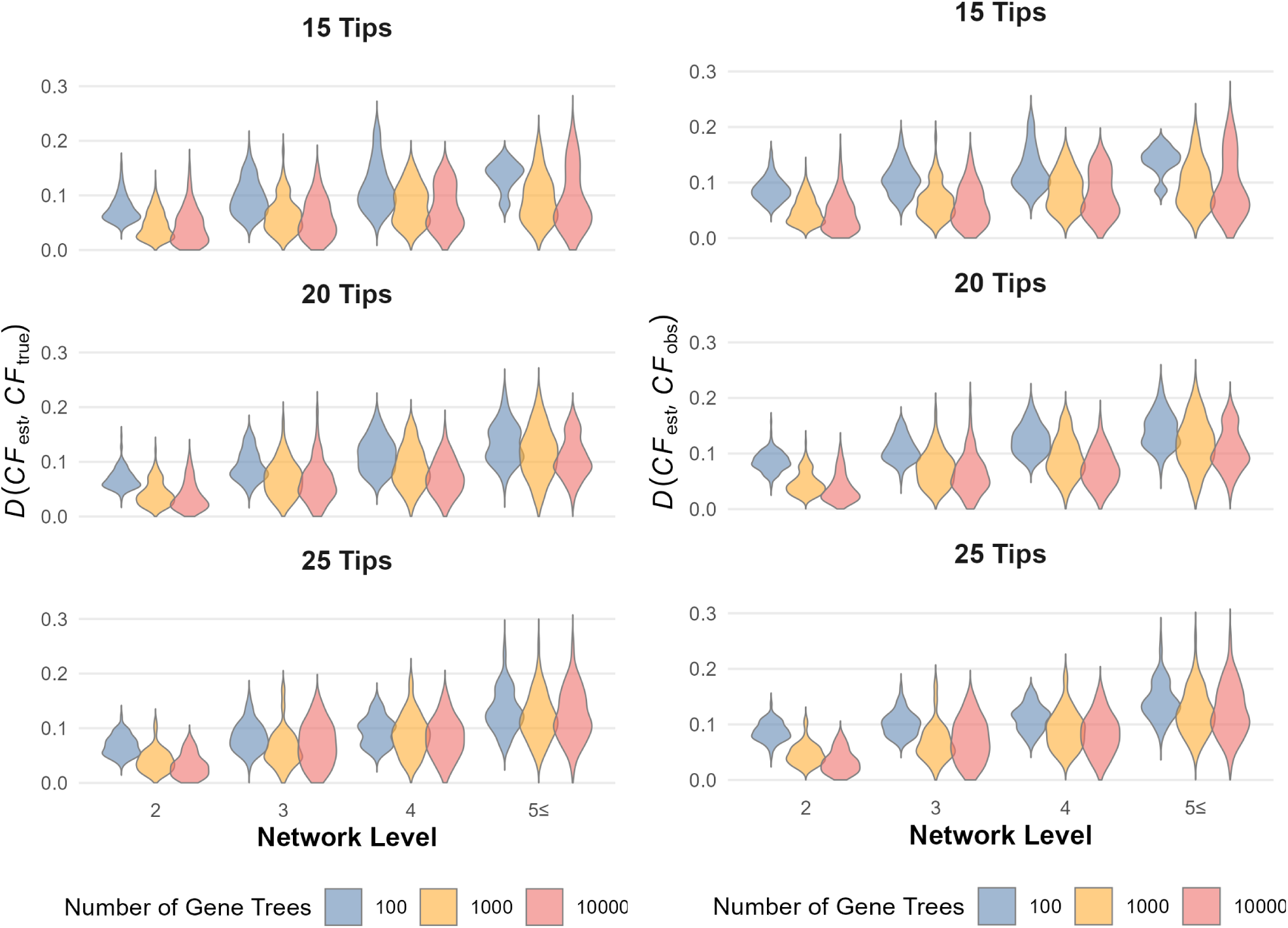
Left: The average distance between the expected CFs from the true network and the expected CFs from the estimated network. Right: The average distance between the observed CFs and the expected CFs from the estimated network.

**Figure S3:**
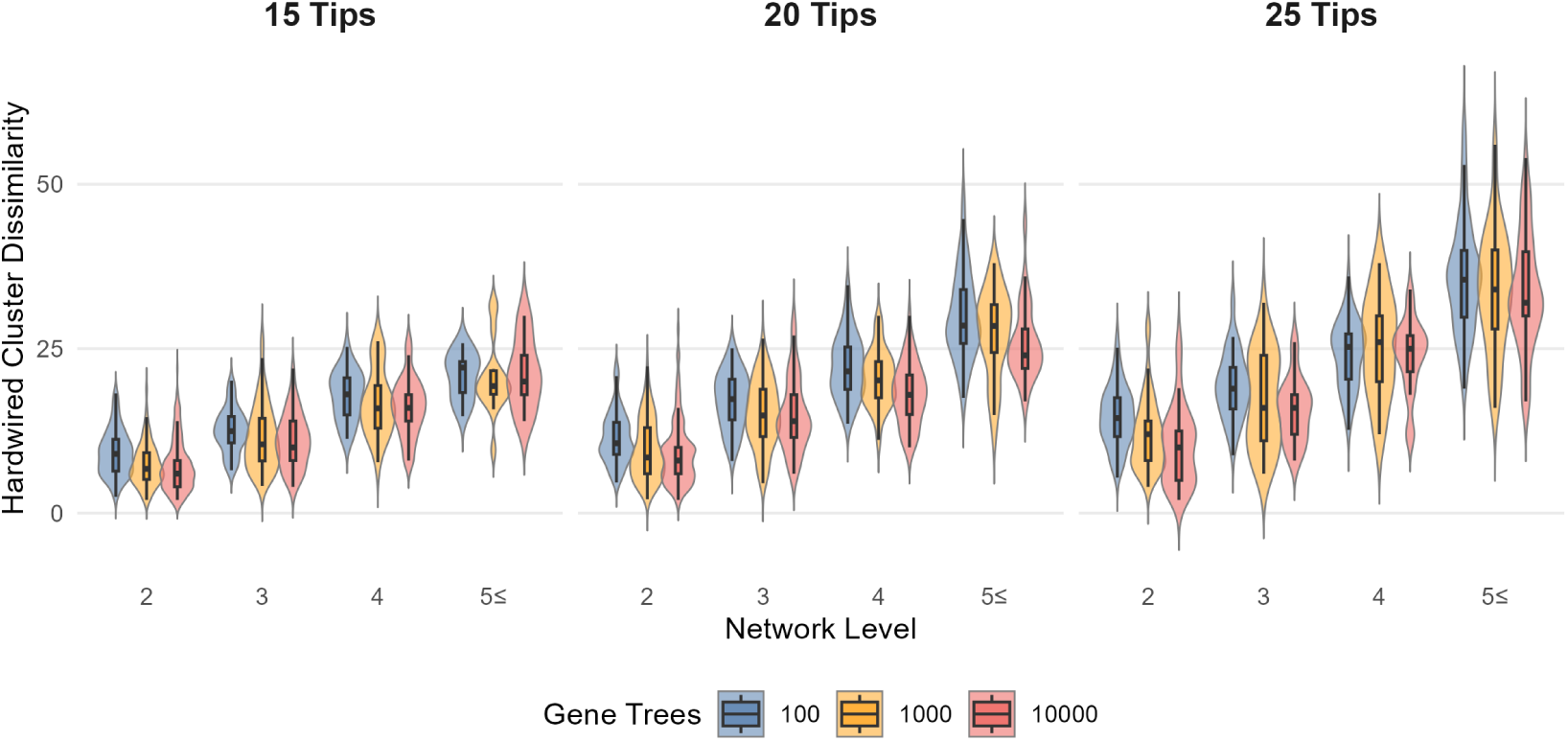
Average Hardwired cluster dissimilarity measure between estimated networks and true networks across replicates when*h*_max_ was set to be the number of blobs in the true network. Each panel summarizes the simulations between both hybridization rates for 15,20, and 25 taxa, respectively. Violin colours denote the number of gene trees used to estimate species networks.

**Figure S4:**
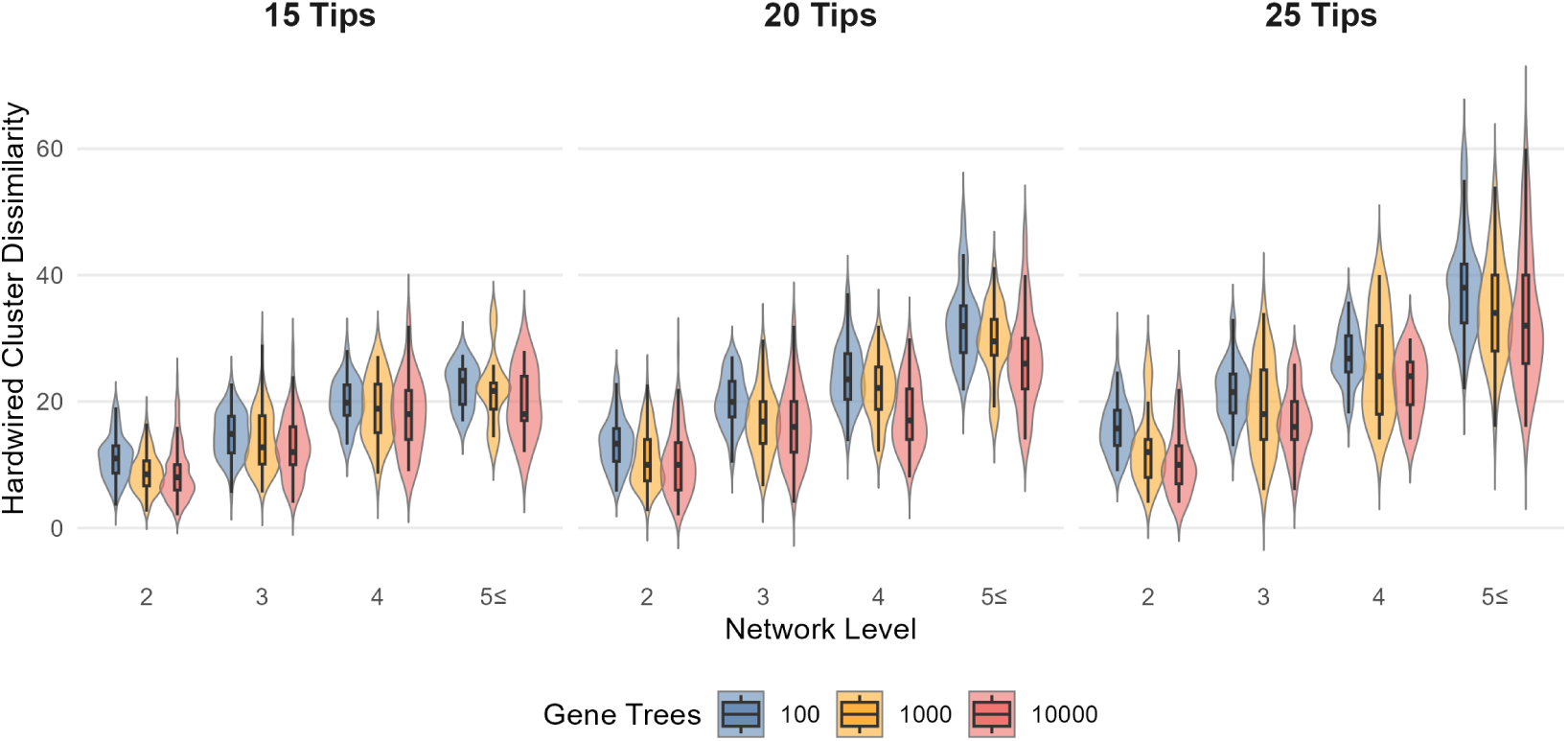
Average HWC dissimilarity. Here *h*_max_ was set to be the number of reticulations in the true network (up to a maximum of 5). Each panel summarizes the simulations between both hybridization rates for 15,20, and 25 taxa, respectively.

**Figure S5:**
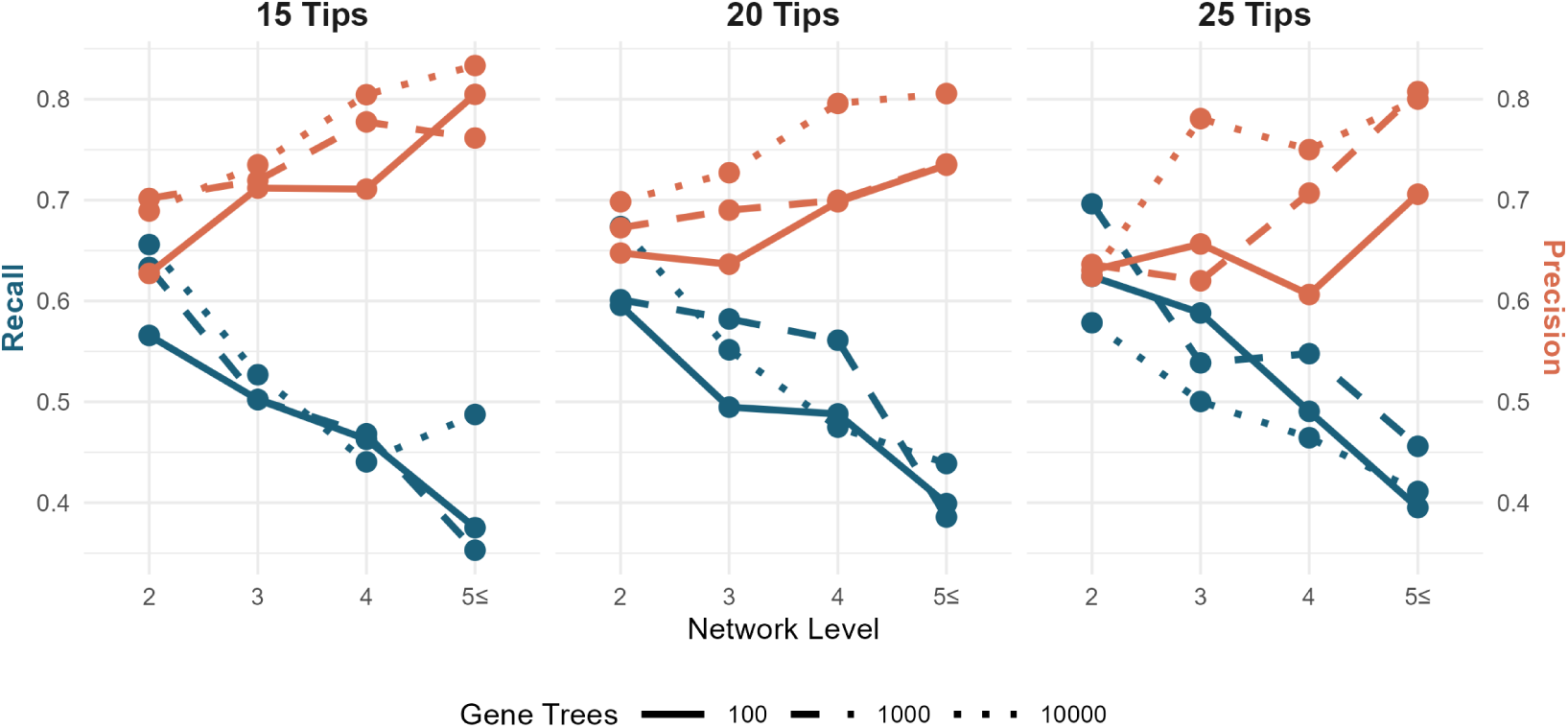
Average recall and precision across replicates for identifying taxa with hybrid descent. Here *h*_max_ was set to be the number of reticulations in the true network (up to a maximum of 5). Each panel summarizes the simulations between both hybridization rates for 15,20, and 25 taxa, respectively.

**Figure S6:**
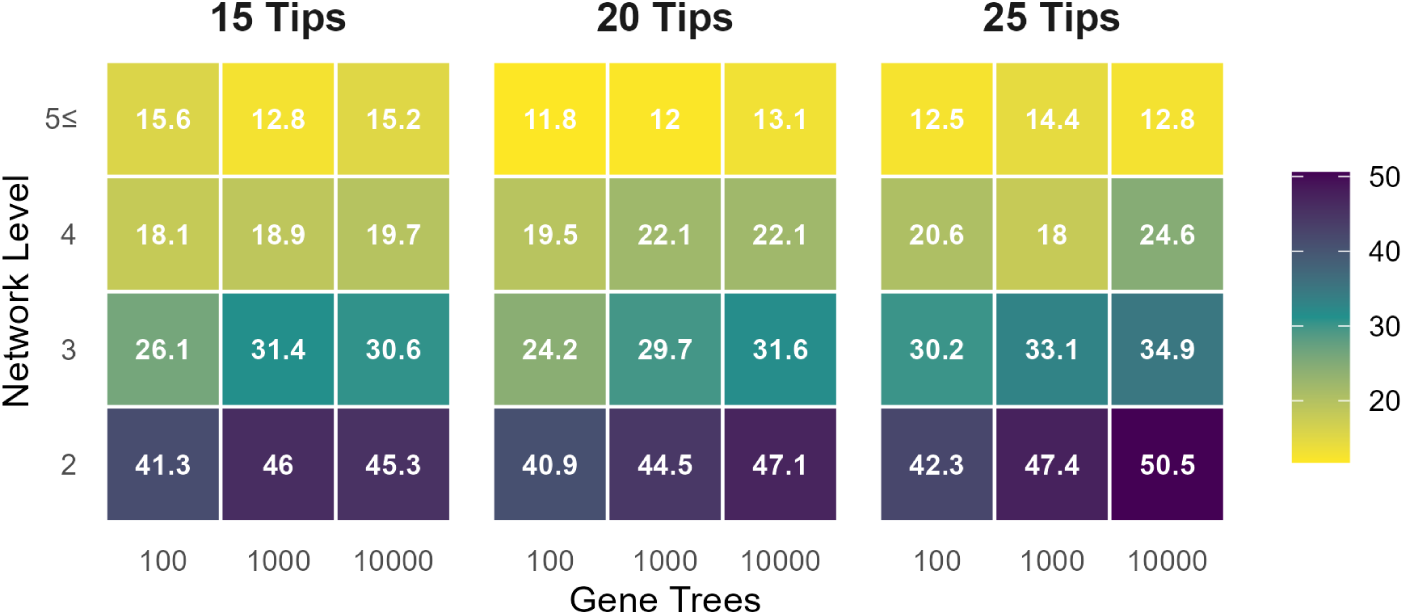
Percentage of true hybrid clades that were exactly found in the estimated network. Exactly found denotes a hybrid node with the same set of descendants as a hybrid node in the true network, not necessarily that the topological relationships in the network are the same between clades.

**Figure S7:**
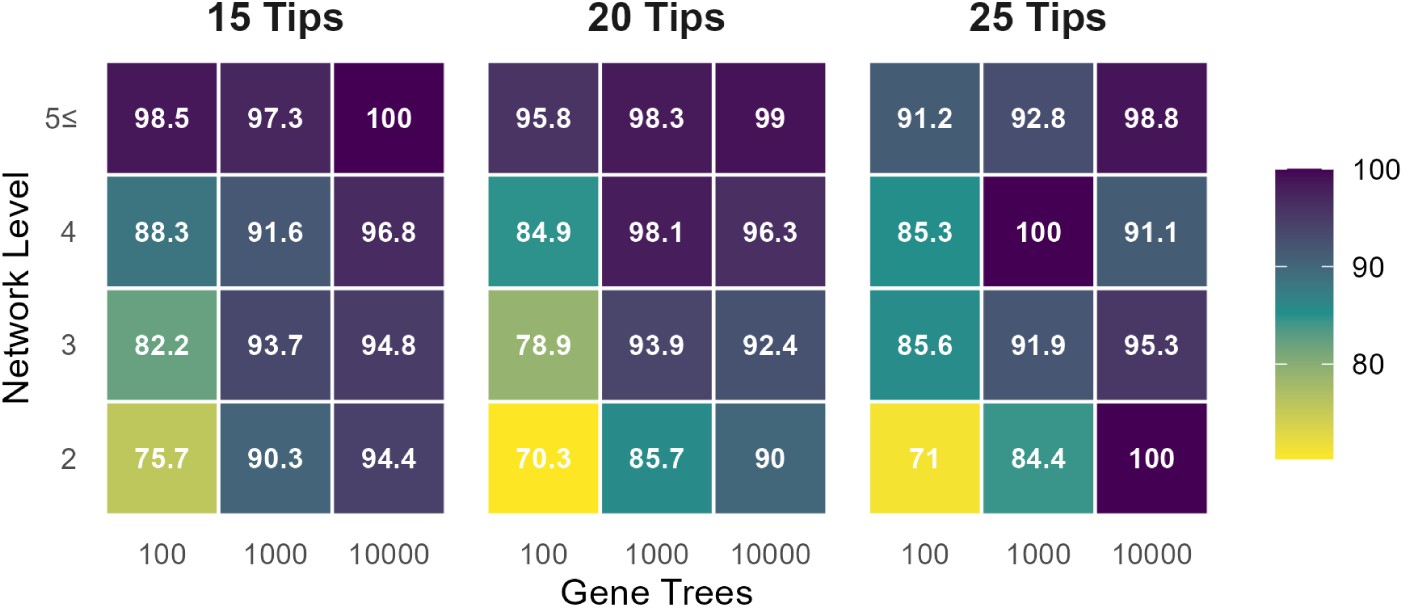
The average proportion of blobs on the true network that were found on estimated networks.

**Figure S8:**
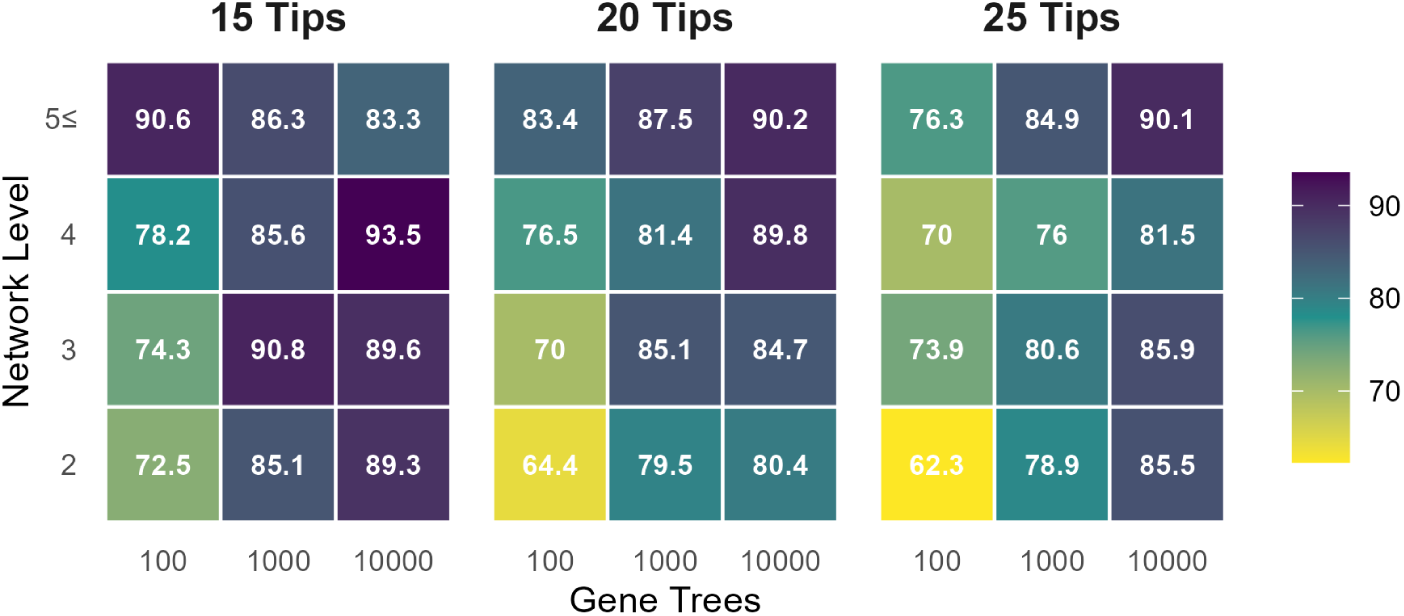
The average proportion of blobs on the estimated network that are compatible with a blob on the true network.

**Figure S9:**
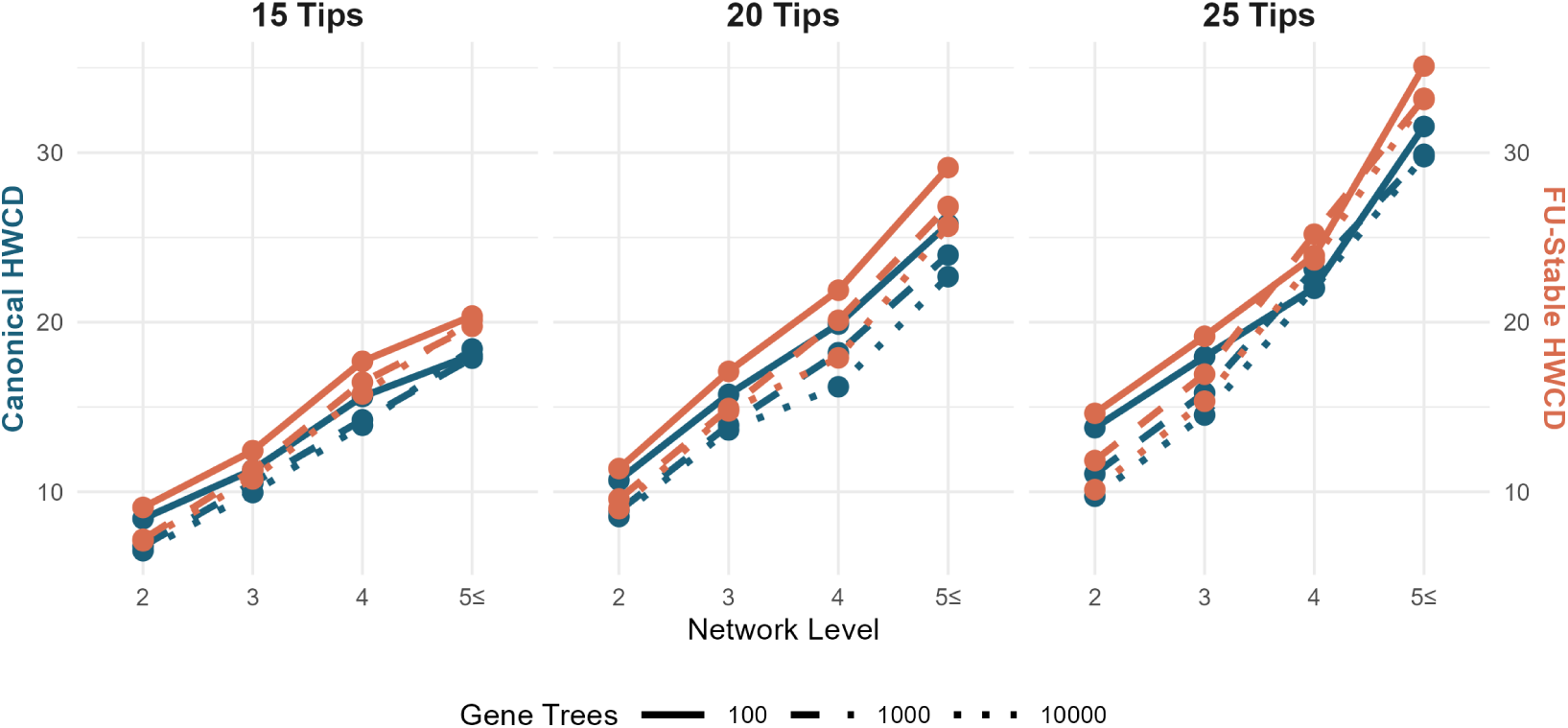
Average HWC dissimilarity scores on reduced form networks. *h*_max_ was set to be the number of blobs in the true network. Each panel summarizes the simulations between both hybridization rates for 15,20, and 25 taxa, respectively.

**Figure S10:**
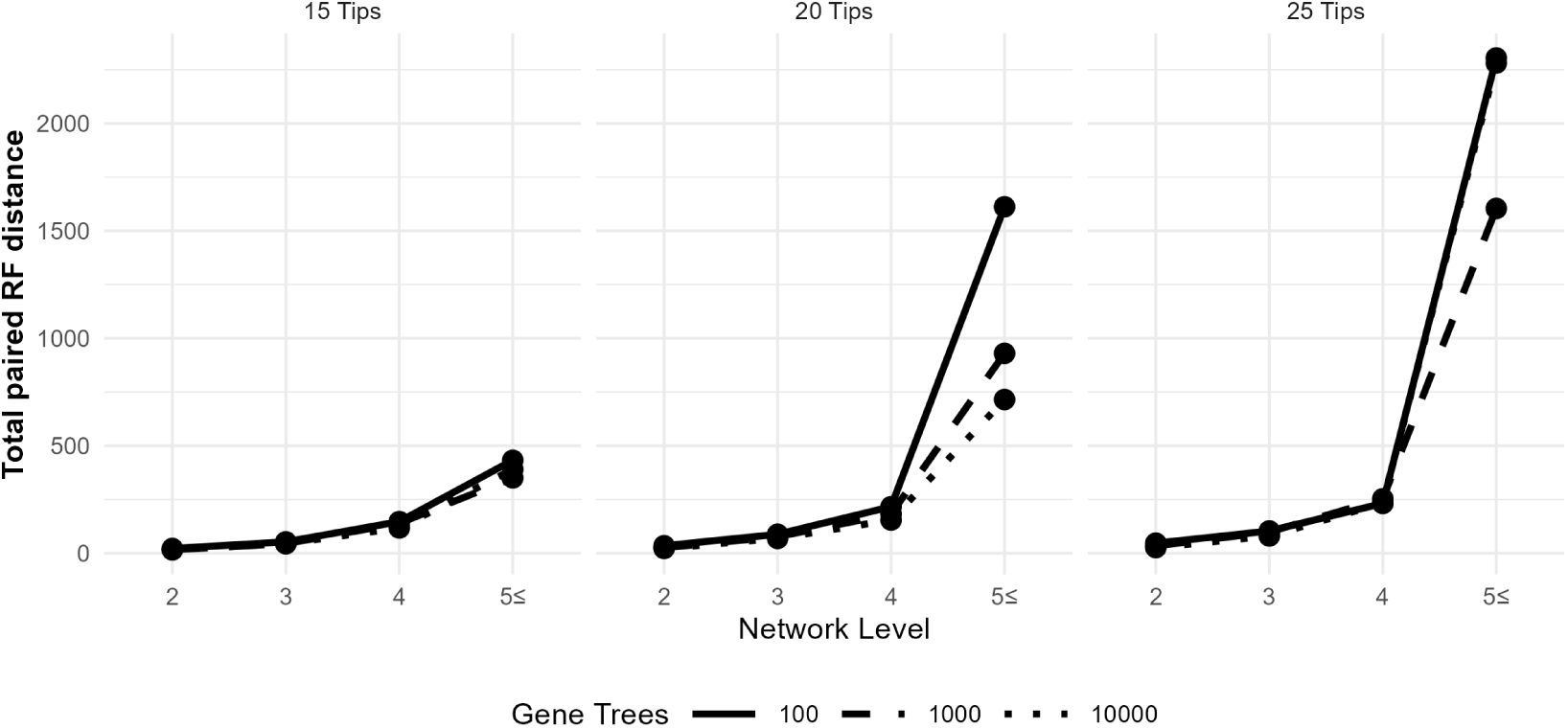
Total distance between true and estimated displayed trees.

**Figure S11:**
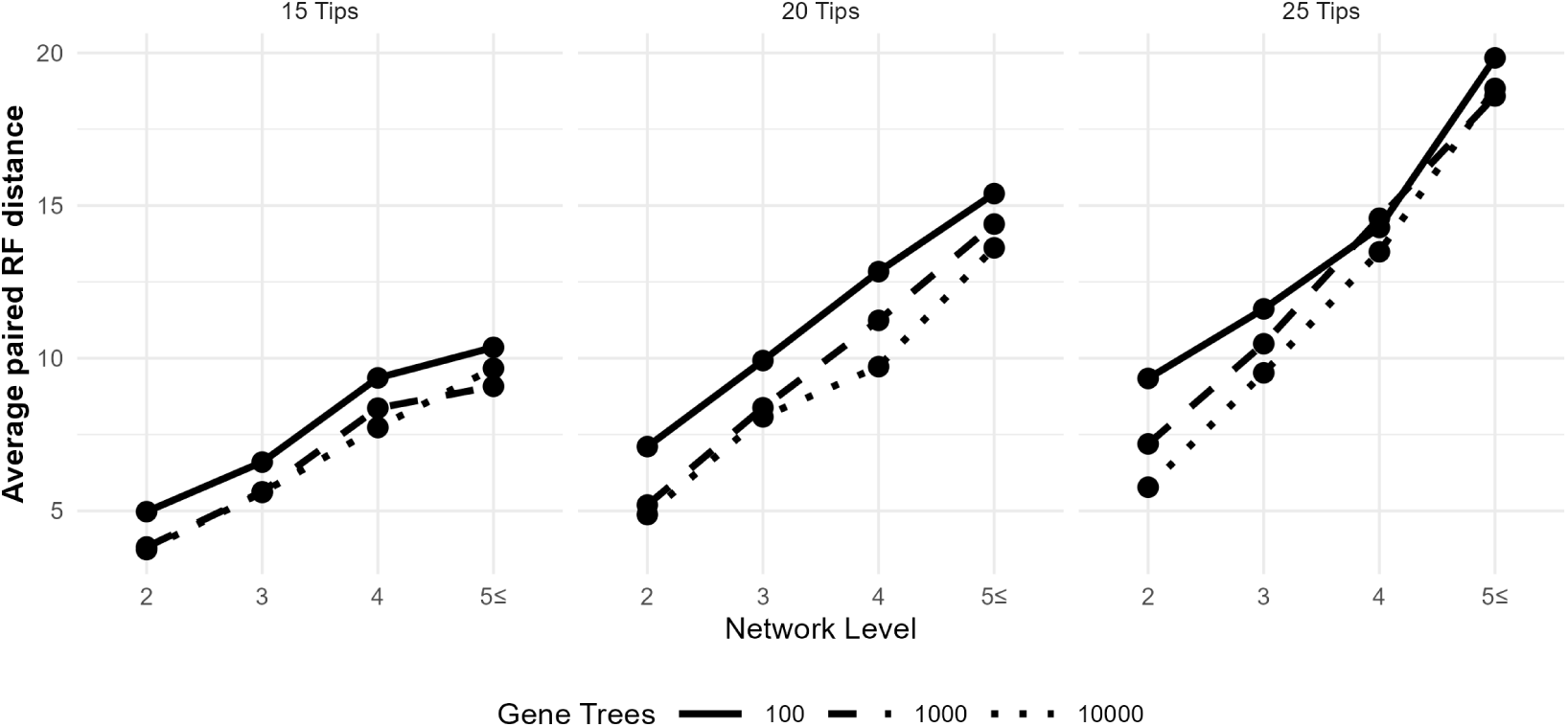
Average distance between true and estimated displayed trees.

